# The interferon-ɣ inducible factor 16 (IFI16) restricts adeno-associated virus serotype 2 (AAV2) transduction in an immune-modulatory independent way

**DOI:** 10.1101/2024.01.22.576730

**Authors:** Sereina O. Sutter, Kurt Tobler, Michael Seyffert, Anouk Lkharrazi, Joël Zöllig, Elisabeth M. Schraner, Bernd Vogt, Hildegard Büning, Cornel Fraefel

## Abstract

We determined the transcription profile of AAV2-infected primary human fibroblasts. Subsequent analysis revealed that cells respond to AAV infection through changes in several significantly affected pathways including cell cycle regulation, chromatin modulation, and innate immune responses. Various assays were performed to validate selected differentially expressed genes and confirmed not only the quality but also the robustness of the raw data. One of the genes upregulated in AAV2 infected cells was the interferon-ɣ inducible factor 16 (IFI16). IFI16 is known as a multifunctional cytosolic and nuclear innate immune sensor for double-stranded, as well as single-stranded DNA, exerting its effects through various mechanisms, such as interferon response, epigenetic modifications, or transcriptional regulation. IFI16 thereby constitutes a restriction factor for many different viruses amongst them, as shown here, AAV2 and thereof derived vectors. Indeed, the post-transcriptional silencing of *IFI16* significantly increased AAV2 transduction efficiency, independent of the structure of the virus/vector genome. We also show that IFI16 exerts its inhibitory effect on AAV2 transduction in an immune-modulatory independent way, by interfering with Sp1-dependent transactivation of wild-type AAV2 and AAV2 vector promoters.

**IMPORTANCE:** Adeno-associated virus (AAV) vectors are among the most frequently used viral vectors for gene therapy. The lack of pathogenicity of the parental virus, the long-term persistence as episomes in non-proliferating cells, and the availability of a variety of AAV serotypes differing in their cellular tropism are advantageous features of this biological nanoparticle. To deepen our understanding of virus-host interactions, especially in terms of innate immune responses, we present here the first transcriptome analysis of AAV serotype 2 (AAV2) infected human primary fibroblasts. Our findings indicate that the interferon-ɣ inducible factor 16 (IFI16) acts as an antiviral factor in AAV2 infection and AAV2 vector-mediated cell transduction in an immune-modulatory independent way by interrupting the Sp1-dependent gene expression from viral or vector genomes.

## INTRODUCTION

Adeno-associated virus serotype 2 (AAV2) is a small, non-pathogenic, helper virus-dependent parvovirus with a single-stranded (ss) DNA genome of approximately 4.7 kb, which has attracted interest as basis for one of the most frequently applied vector system in human gene therapy (1). In absence of a helper virus, AAV2 can integrate its genome site-preferentially into the adeno-associated virus pre-integration site (AAVS1) on human chromosome 19 or persist in an episomal form in the nucleus (2, 3). Co-infection with a helper virus, such as herpes simplex virus type 1 (HSV-1), leads to the entrance into a lytic replication cycle including the production of progeny virus particles (4). The AAV2 genome consists of two large open reading frames (ORFs) flanked by 145 nt long inverted terminal repeats (ITRs) located on either side. The *rep* gene encodes the four non-structural Rep proteins, two of which are transcribed from the p5 and the p19 promoter, respectively. An alternative splice site at map position 42 and 46 regulates the expression of the alternative transcripts (5, 6). The four different Rep proteins are designated according to their molecular weight as Rep78, Rep68, Rep52 and Rep40. The promoter activity is regulated by the Rep binding site (RBS), which allows the Rep protein to act as either a transactivator or repressor (7). In the absence of a helper virus, only little synthesis of Rep proteins takes place, which nonetheless is sufficient to repress any further transcription. The three structural proteins VP1, VP2 and VP3, building up the icosahedral capsid, are encoded by the *cap* gene. Moreover, the *cap* gene encodes the assembly-activating protein (AAP) and the membrane-associated accessory protein (MAAP) by means of a nested alternative ORF (8, 9).

RNA sequencing (RNA-seq) is a technology which uses the commitment of next generation sequencing (NGS), also known as deep sequencing, to identify transcripts and their quantity in cells at a given time point (10). The development of NGS with its high base coverage and sample throughput facilitates the sequencing of transcripts in cells and allows to study alternative spliced transcripts, changes in gene expression and cellular pathway alterations during infection (11). Moreover, RNA-seq facilitates a closer look at different types of RNA (e.g. mRNA, sRNA, tRNA, miRNA), ribosome profiling, and the total RNA content of a cell (12) by overcoming the limited coverage and inability to detect rare transcript variants. Using this approach, we provide here a genome-wide expression profile of AAV2-infected cultured fibroblasts, unveiling virus-host interactions. Transcript mapping, followed by an overall expression counting resulted in 44‘175 annotations. The deeper analysis of differently expressed genes between AAV2- and mock-infected cells revealed 1‘929 distinct (p < 0.01, number of reads ≥ 40) regulated genes of which 92.78% were protein coding, including amongst others the interferon-inducible p200-family protein IFI16. IFI16 is assumed to be an innate immune sensor for cytosolic and nuclear double-stranded (ds), as well as ssDNA (13). IFI16 has been shown to be a restriction factor for many different viruses through various mechanisms, including interferon response, transcriptional regulation, and epigenetic modifications. For example, human cytomegalovirus (HCMV) replication was shown to be significantly enhanced due to IFI16-mediated blockage of Sp1-dependent transcription of UL54 (14). Moreover, IFI16 can also restrict HSV-1 replication by repressing HSV-1 gene expression, independently of its roles in the immune response (13), via global histone modifications by decreasing the markers for active chromatin and increasing the markers for repressive chromatin on cellular and viral genes (15, 16). IFI16, however, inhibits not only various DNA viruses such as HCMV, HSV-1 or human papillomavirus 18 (HPV18) (17) but shares properties of known anti-retroviral restriction factors (18) and blocks human immunodeficiency virus type 1 (HIV-1) by binding and inhibiting the host transcription factor Sp1 that drives viral gene expression (19), similar to HCMV.

AAV2 and AAV2 vectors deliver ssDNA or dsDNA or induce the formation of ss-, ds-, and circular dsDNA products and may therefore provoke an IFI16 triggered reaction. Consequently, we aimed to address the question whether IFI16 influences AAV2 infection and vector-mediated cell transduction, respectively.

## RESULTS

### Total RNA-seq reveals 1‘929 differentially expressed genes in AAV2-infected versus mock-infected normal human fibroblasts

Structural cells, such as fibroblasts, are found in literally every tissue, making them susceptible to a variety of AAV serotypes and thereof derived vectors. Although their molecular signature is not maintained between organs (20), tissue resident fibroblasts were shown to play a key role in the suppression or activation of immune responses (reviewed in (21)). To assess the global gene expression profile of AAV2 and mock-infected normal human fibroblast (NHF) cells, RNA-seq was performed on total RNA isolated 24 hours post infection (hpi). Transcript mapping to the human assembly and gene annotation from Ensembl, extended by the AAV2 sequence, followed by an overall expression counting (see Materials and Methods), resulted in 44‘175 annotations. The analysis revealed 1‘929 differentially expressed (DE) genes (p < 0.01, number of reads ≥ 40) between AAV2 and mock-infected cells of which 92.78% were protein coding. A small portion of the restricted annotation resulted in pseudo genes and anti-sense RNA (Table 1).

**TABLE 1.** Summary of the gene annotation and numbers of differentially expressed genes at 24 hpi.

| Type | Annotation | p < 0.01, number of reads ≥ 40 |
| --- | --- | --- |
| Protein coding | 19`934 | 1`787 |
| Small RNA | 1`286 | 2 |
| rRNA | 102 | - |
| Pseudogene | 9`948 | 70 |
| lincRNA | 5`862 | 13 |
| Antisense | 4`701 | 41 |
| Other | 2`339 | 13 |
| AAV specific | 3 | 3 |
| Total | 44`175 | 1`929 |

### The biological process ontology of AAV2-infected cells

To further explore the 1‘929 differentially regulated genes, a Gene Ontology (GO) term biological process (BP) analysis was performed. The GOterms BP were identified by DAVID (22, 23) and graphically visualized as enrichment map using Cytoscape. Eight distinct clusters of biological processes which differed between AAV2 and mock-infected cells became evident (Fig. 1). The cluster termed “regulation of macromolecule / metabolic processes / gene expression” was heavily represented with 38 nodes, followed by the cluster “cell cycle regulation” with 30 nodes. Considering that the size of the nodes within the clusters corresponded to the number of genes included, some GOterms were represented by more genes than others. Taking this into account, the clusters “cell cycle regulation”, “chromatin organization”, “DNA replication / damage response” and “apoptosis” were highly represented. This suggested that AAV2 can modulate crossroads of host gene expression relevant for cell cycle regulation, chromatin modulation, DNA-damage response (DDR) and apoptosis. To further analyze and visualize the DE genes within the cell cycle and chromatin organization clusters, unsupervised hierarchical clustering of the expression levels from the 50 most differently expressed genes of those GOterms was generated. The DE genes were ordered according to their absolute difference in expression with the most differentially regulated genes at the top (see Fig. S1 in the supplemental material). The highest fold difference in the chromatin organization cluster was 5.4 for *HIST1H2AJ*, the smallest difference of 2.2 for *MCM2*, whereby the majority of differentially expressed genes belonged to the histone cluster 1 genes. In the case of the cell cycle regulation cluster, *CDC45* and *UBE2I* showed a maximum/minimum fold difference of 2.8 and 1.8, respectively. Overall, the differential expression profile of the cell cycle showed a broader range of genes included, compared to the uniform pattern of the chromatin organization group, consisting mostly of histone cluster 1 genes.

**FIG 1.**
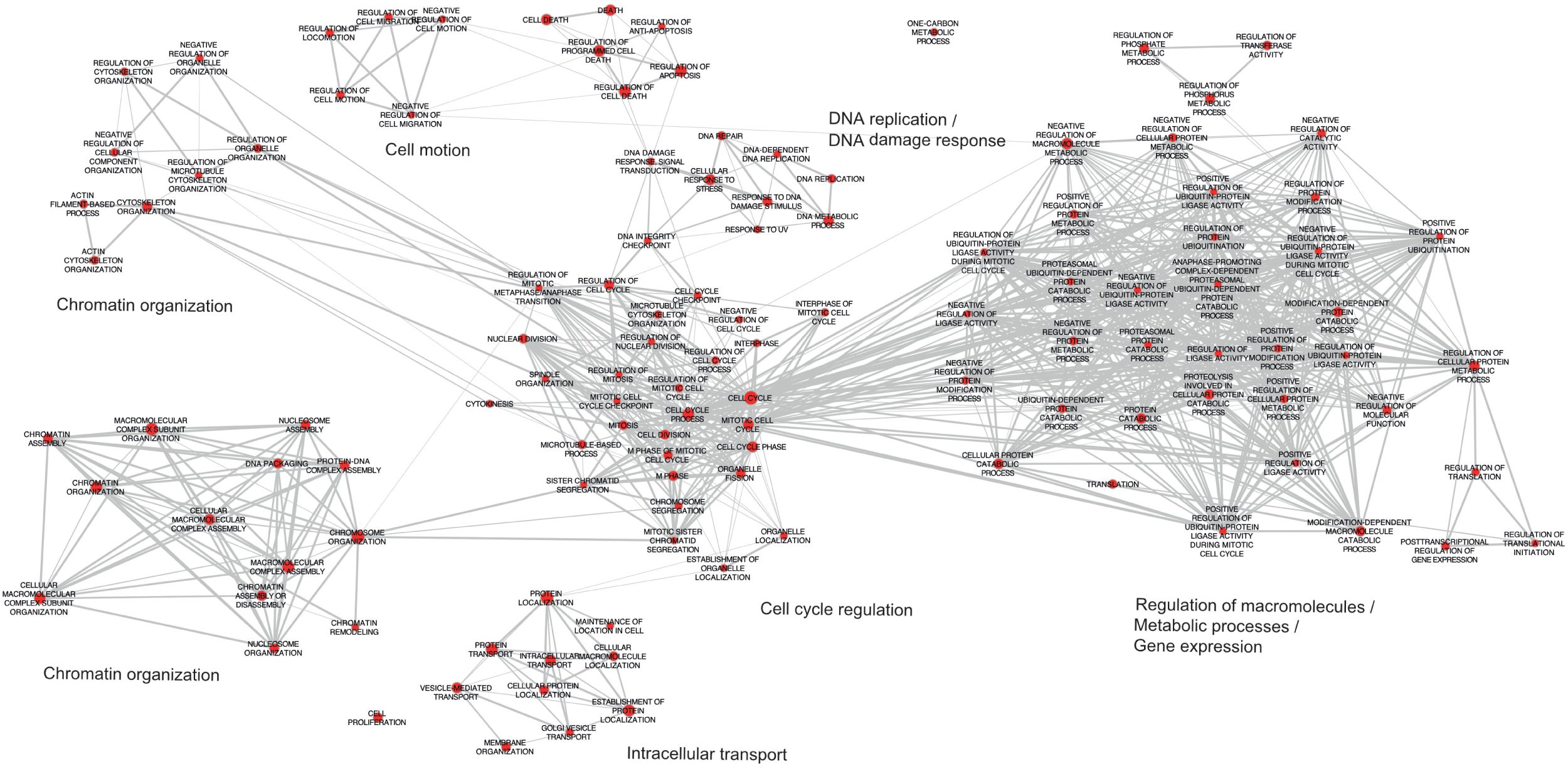
Enrichment map of the DAVID GOterms. Analysis of 1930 differentially expressed genes (p < 0.01, number of reads > 40) by DAVID was used to create an enrichment map of the GOterms using Cytoscape. Nodes (red) represent the individual GOterms, while their size corresponds to the number of genes included. Edges (grey lines) represent mutual overlaps and the thickness the number of overlaps. The most affected biological processes are summarized as keywords.

### Identification of the number of genes with a fold change of ≥ 1.5

The remaining 1‘929 DE genes were further screened with more restrictive criteria. Specifically, genes with a fold change (FC) higher than 1.5, which is equivalent to a log_2_ ratio of the mean transcript abundance of AAV2 to mock-infected cells of > |0.58|, were selected. Overall, 268 (Fig. 2A) and 604 (Fig. 2C) cellular genes were down- or upregulated, respectively. Figure 2B, however, illustrates the entire distribution frequency of the log_2_ ratio of the mean read abundance of the transcripts of AAV2 to mock-infected cells. Overall, 872 genes were defined as differentially expressed at least 1.5-fold in AAV2-infected cells compared to mock-infected cells. To compare the transcript read abundance of the 872 genes between AAV2-infected and mock-infected cells, an unsupervised clustered heat map was generated (see Fig. S2 in the supplemental material). The read abundance of transcripts (log_2_) ranged from roughly 0.54 (M2) to approximately 16 (A3). Further, the heat map showed that the 3 samples from AAV2-infected cells were homogeneous and clearly differed from the mock-infected cells. However, the mock-infected M2 sample did not correlate with M1 and M3.

**FIG 2.**
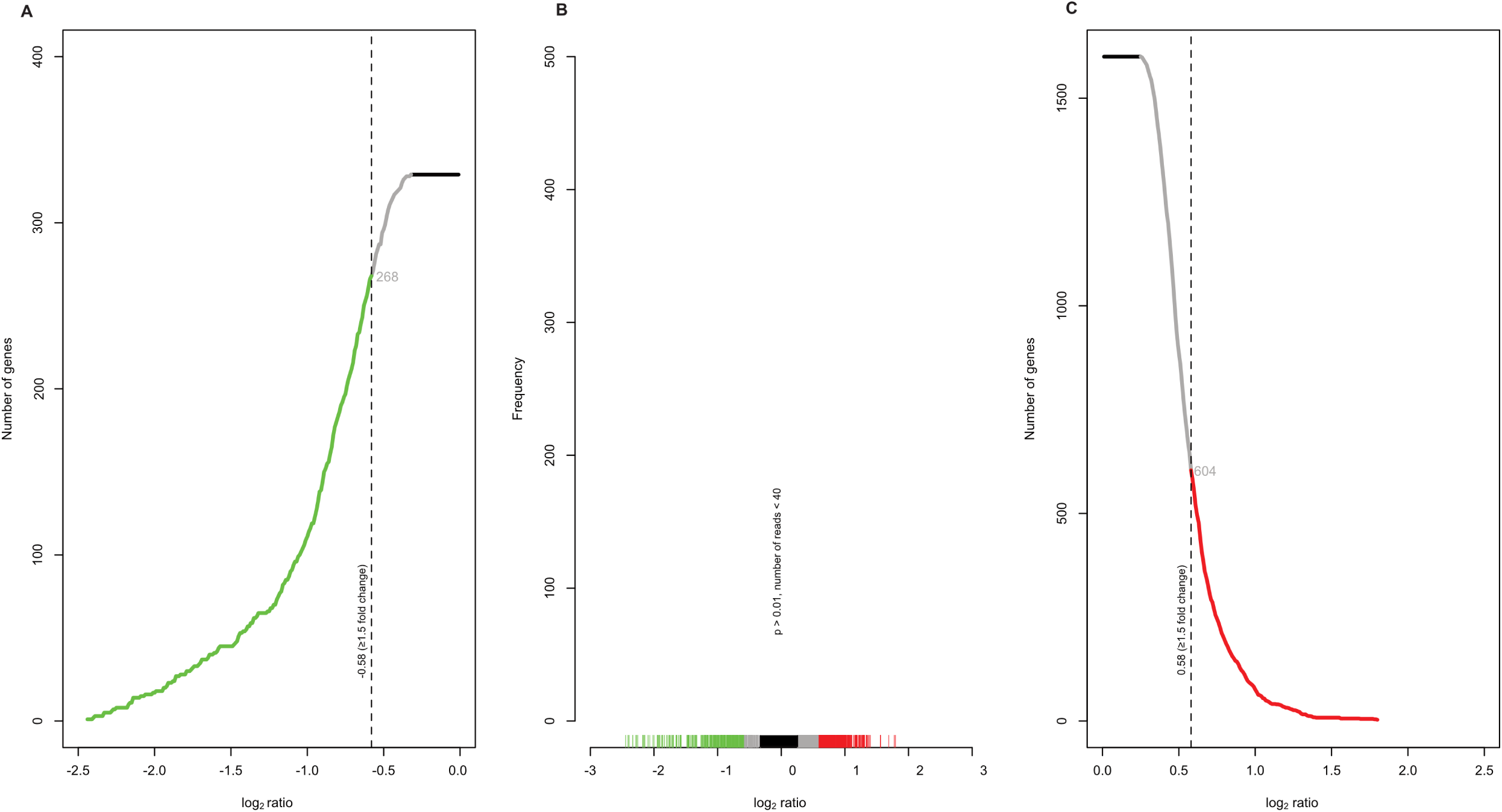
Identification of the number of genes with a fold change of ≥ 1.5. The number of genes with a log_2_ ratio of mean read abundance of transcripts (AAV2-infected NHF cells to mock-infected NHF cells) between (A) -0.01 and -2.5 or (C) 0.01 and 2.5, respectively and a significance threshold of p < 0.01. The dashed vertical line in -0.58 and (C) 0.58 indicates a fold change of 1.5. The number of genes representing a fold change ≥ 1.5 are numerically indicated in grey (872 genes in total). The number of genes (A) below a log_2_ ratio of -0.58 are illustrated in green, whereas those (C) with a log_2_ ratio greater than 0.58 are depicted in red. The same color code was used for (B) the distribution frequency. The black bar in (B) indicates those log_2_ values, which were directly excluded.

### Functional classification of the transcriptome following AAV2 infection

To further analyze the RNA-seq data, the list of 872 differentially expressed genes with an FC ≥ 1.5 was projected onto KEGG pathways (24). The KEGG analysis revealed that the majority of the genes involved in cell cycle regulation were downregulated upon AAV2 infection (Fig. 3). Several cyclins as well as their binding partners, the cyclin-dependent kinases (CDKs), which are relevant for the progression of the cell into S-phase and further transition into G2-phase, were downregulated. These downregulations also influence the expression state of several transcription factors (E2Fs), spindle checkpoint proteins (MAD2, BUBR1) and replication relevant proteins (MCMs, ORC). Moreover, some anaphase-promoting complex (APC) proteins (ESP1, PTTG) were also downregulated. Some of the upregulated genes negatively regulate those genes that were downregulated, such as the growth arrest and DNA-damage-inducible protein (GADD45), which negatively affects the binding of CDK1 to cyclin B1 (25, 26), and the CDK inhibitory protein (CIP1), also known as CDKN1A or p21. Others, like the 14-3-3 σ protein (encoded by *YWHAB*), D-type cyclins and the oncoprotein MDM2 have a direct influence on the cell cycle progression. The upregulated abelson tyrosine-protein kinase 1 (ABL1) is involved in DNA-damage response and apoptosis, as well as in the phosphorylation of several cell cycle relevant proteins, such as the retinoblastoma (RB) protein (27) or proteasome subunit alpha type-7 (PSMA7), thereby influencing their activation state and protein interactions.

**FIG 3.**
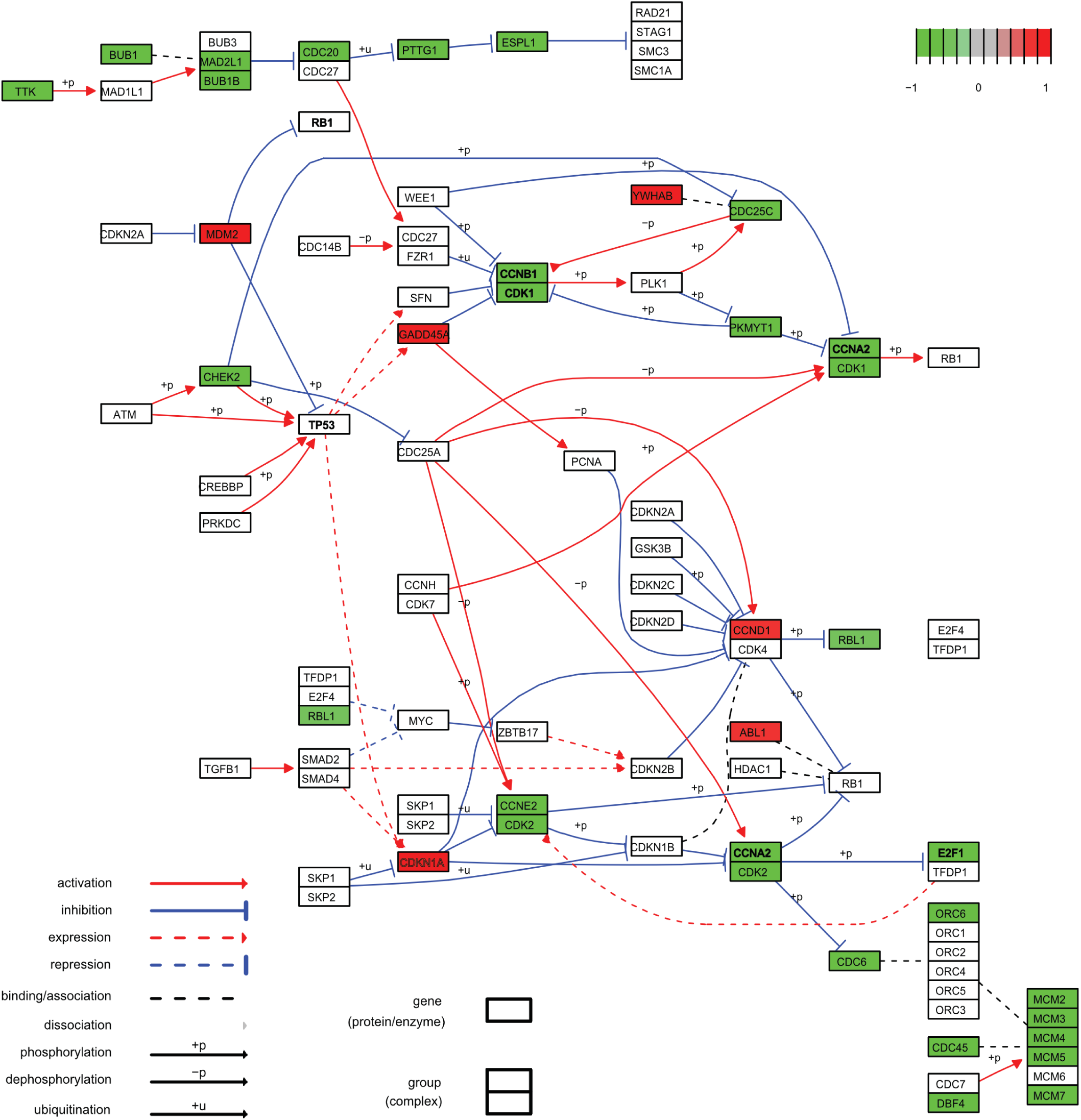
KEGG pathway analysis of the cell cycle allowed the identification of differentially expressed genes in AAV2 and mock-infected cells. Upregulated genes are color coded in red, while downregulated genes are depicted in green (FC ≥ 1.5, p < 0.01, number of reads > 40). Symbol legend is shown in the KEGG pathway analysis.

### Validation of RNA-seq data on transcription and protein level

In order to confirm the observations gained by the RNA-seq analysis, transcript and protein levels of selected genes (Fig. 3, in bold) were evaluated by quantitative reverse transcription PCR (RT-qPCR) and Western blot analysis, respectively (Fig. 4). However, due to the heterogeneous nature of the M2 sample, the selection of genes for validation on transcript and protein level was based only on the RNAseq data obtained from the M1 and M3 samples. The RT-qPCR data was normalized to the mean value of the transcriptional activity of two housekeeping genes (*GAPDH* and *SDHA*), since their expression profile was only marginally affected (SDHA; log_2_ = 0.2, p = 0.3 GAPDH; log_2_ = 0.2, p = 0.04) upon AAV2 infection. The resulting ΔΔC_t_ values of each gene of interest were plotted against their corresponding log_2_ ratio of the mean transcript abundance of AAV2 to mock-infected cells from the RNA-seq data (Fig. 4A). The expression levels of *CCNA2*, *CCNB1*, *CDK1* and *RB1* correlated well between the RT-qPCR and the RNA-seq data (R^2^ = 0.897), although for *E2F1*, *CDKN1A* and *TP53* a moderate difference of the RT-qPCR data to the RNA-seq data was observed. Overall, the results from the RNA-seq analysis were confirmed by RT-qPCR.

**FIG 4.**
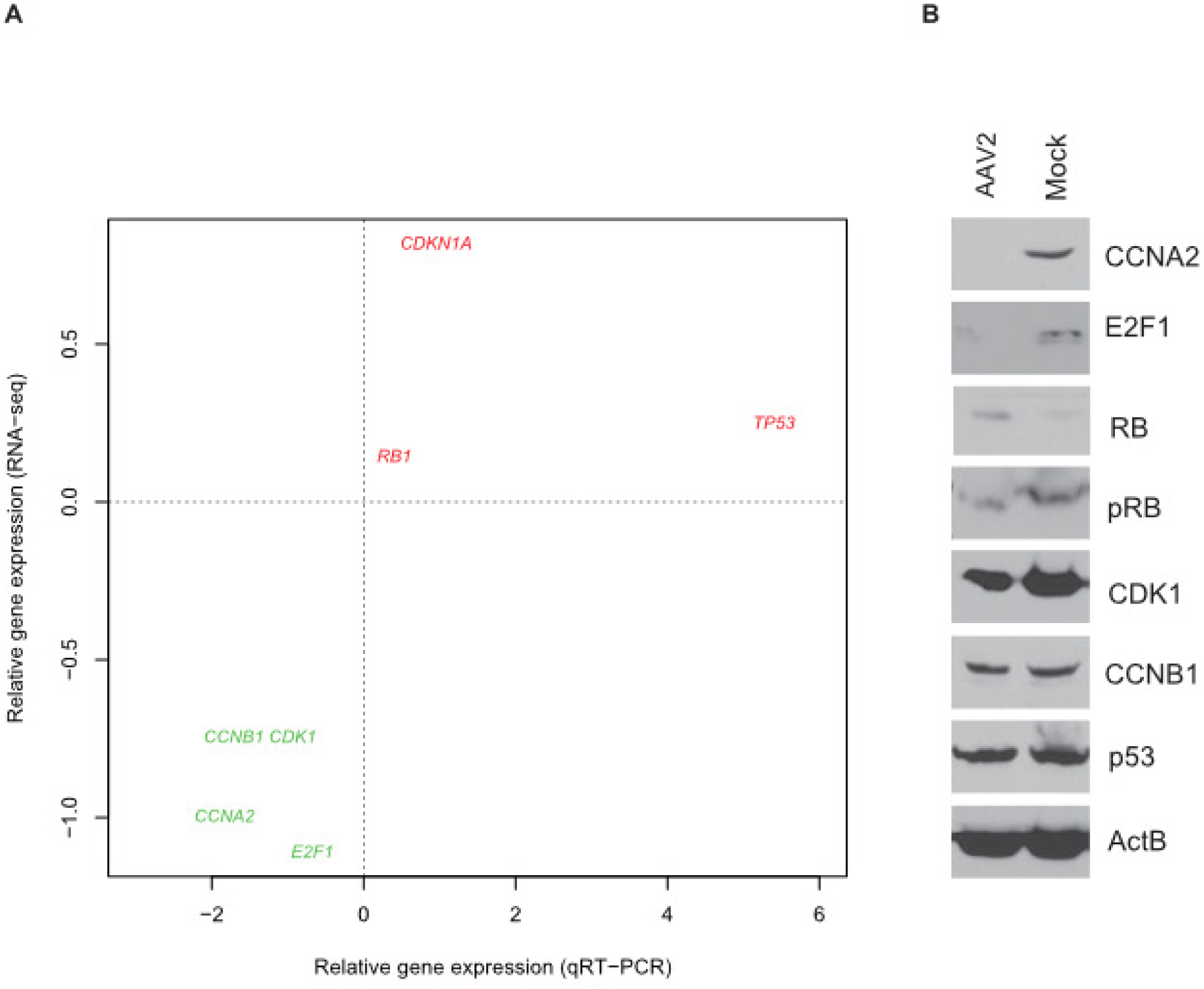
Validation of RNA-seq gene expression profile by RT-qPCR and Western blot analysis. (A) Comparison of relative gene expression profile of selected genes from AAV2-infected versus mock-infected NHF cells at 24 hpi by RT-qPCR and RNA-seq. Upregulated genes are color coded in red while downregulated genes are depicted in green. All data generated by RT-qPCR were normalized against the mean of two housekeeping genes *GAPDH* and *SDHA*. Data shown are the means of triplicates. (B) Total cell lysates of mock-infected (Mock) and AAV2-infected cells prepared at 48 hpi were subjected to Western blot analysis. The β-actin stain was performed as a loading control.

Western blot analysis of total cell lysates at 48 hpi was performed to evaluate the protein expression levels of selected genes recognized by RNA-seq to be differentially expressed between AAV2-infected and mock-infected cells. The 48-h time point of AAV2 infection was chosen because the differential regulation on protein level might lag behind that observed on the level of transcription. The immunoblotting, using specific antibodies for each protein of interest (Fig. 4B), showed a strong reduction of E2F1, CCNA2 and CDK1 on protein level in AAV2-infected cells. No obvious change after the AAV2 infection was detected for total CCNB1 levels, whereas a moderate increase in total RB levels was observed and almost no change in total p53 levels, showing that the RNA-seq and total protein levels accorded well. To further assess the downregulation of E2F1 in AAV2-infected cells, the level of activated RB protein was determined by Western blot analysis using a phospho-specific antibody. Since E2F1 is kept in an inactive state through its interaction with the hypophosphorylated form of RB, enhanced activity of cyclin E and A would normally lead to the phosphorylation of RB, resulting in a reduced affinity of RB for E2F1, thereby causing its release and activation, leading to enhanced expression of cyclin A and other S-phase genes. This protein interplay between cyclin A, E2F1 and the activation state of RB can be readily observed in Fig. 4B (mock-infected cells). In contrast, a reduced level of activated RB, E2F1, CCNA2 and elevated level of hypophosphorylated RB in the AAV2-infected cells was observed. Overall, protein expression levels of the selected genes were in agreement with the transcription levels and connote a differential regulation of genes relevant for cell cycle progression upon AAV2 infection.

### Effect of different AAV2 vectors on the expression profile of selected genes in cell cycle regulation

To address the question whether the single-stranded AAV2 DNA *per se* or low level AAV2 *rep* gene expression, as *rep* transcripts were indeed identified by RNA-seq, caused the changes in host gene expression, NHF cells were infected with AAV2, UV-irradiated AAV2 (UV-AAV2), or a AAV2 vector delivering a vector genome in the self-complementary (sc) genome configuration expressing an enhanced green fluorescent protein (eGFP; scAAVeGFP). The expression of four genes, selected on the basis of their expression profile in RNA-seq/RT-qPCR and relevance in cell cycle regulation, was measured by RT-qPCR. To this end, the RT-qPCR data was normalized to the activity of the housekeeping gene *SDHA*, whose expression was not significantly different between the samples, and the relative expression changes were expressed as ΔΔC_t_ value. As shown in Fig. 5, the four tested genes (*CCNA2*, *CCNB1*, *CDKN1A* and *TP53*) showed remarkable differences in their expression profile between AAV2-infected cells and cells infected with scAAVeGFP. In contrast to this, the expression profiles of *CCNA2*, *CCNB1* and *TP53* did not show a prominent difference between AAV2 and UV-AAV2. Only the relative expression of *CDKN1A*, encoding p21, differed between AAV2 and UV-AAV2 as well as scAAVeGFP. p21 is a key regulator able to arrest cells in G1 and G2 phases. Moreover, *CDKN1A* is known to be induced through the transcriptional stimulation of p53. Increased levels of p21, in turn, are able to inhibit the activity of cyclin A-CDK2, thereby preventing cell cycle progression in G1/S and G2/M. *CCNB1*, in contrast, is predominantly expressed in G2/M phase of the cell cycle, since its gene product is relevant for the formation of the maturation-promoting factor (MPF). p53 is known to induce a cell cycle arrest in G1 phase in response to DDR (28), but additionally p53 has been associated with cell cycle arrest in G2 phase via the induction of *14-3-3 σ* gene expression (29). The results show, that the expression profiles of the selected genes were similar between cells infected with AAV2 or UV-AAV2 and differed from those infected with scAAVeGFP. The expression profiles of AAV2 and UV-AAV2-infected cells implied a cell cycle arrest upon infection while that of scAAVeGFP infected cells did not. This finding is in accordance with previous observations, showing that infection with scAAV2 vectors allowed the cells to progress through mitosis, an event that occurred significantly less frequently upon infection with a single-stranded AAV2 vector (rAAV2) (30).

**FIG 5.**
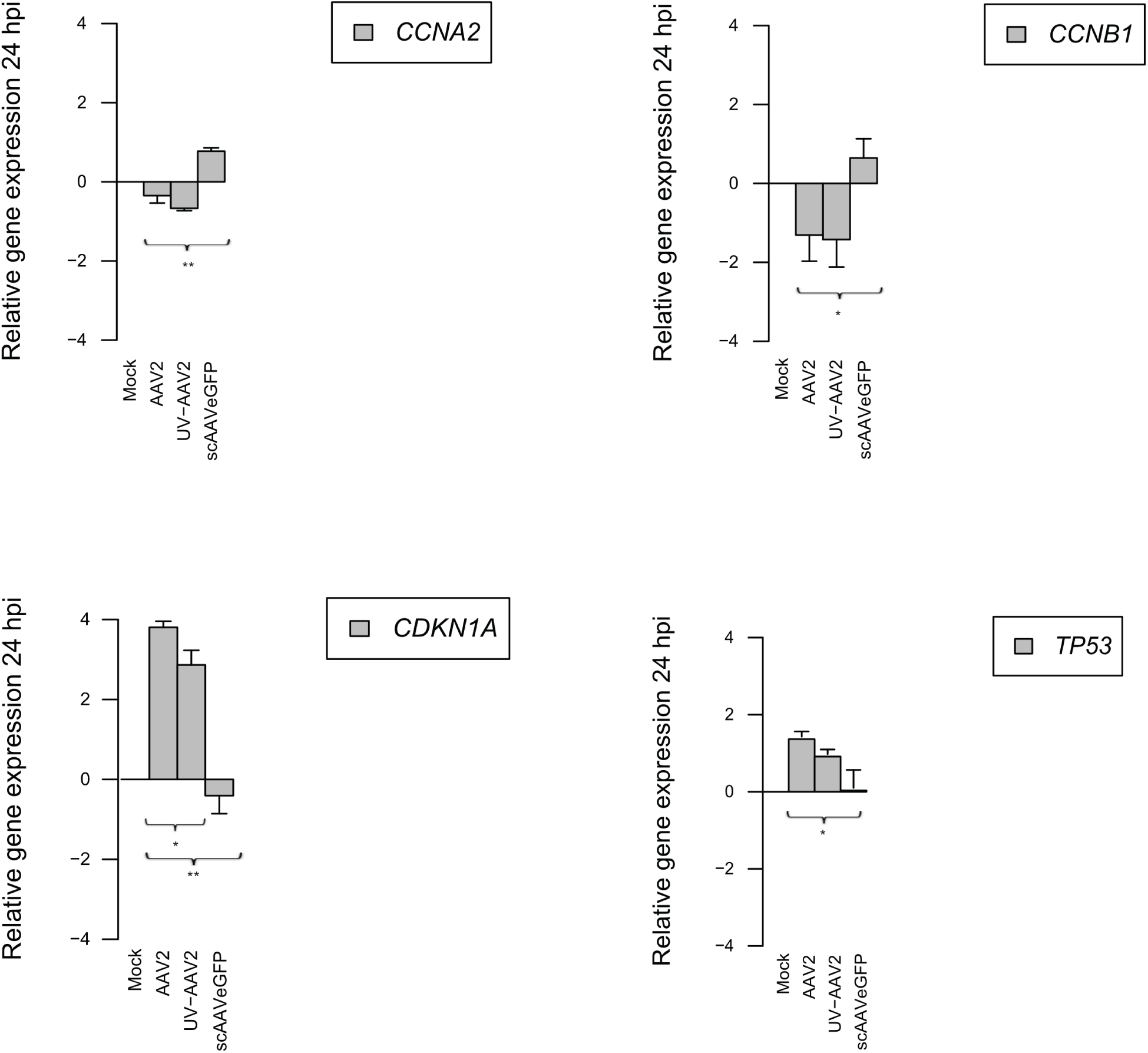
Expression profiles of selected genes involved in cell cycle regulation. Relative gene expression profiles of selected genes in NHF cells infected with AAV2, UV-irradiated AAV2 (UV-AAV2) or self-complementary AAV2 (scAAVeGFP) at 24 hpi. All data generated by RT-qPCR was normalized against the housekeeping gene *SDHA*. Graphs show mean and standard deviation (SD) from triplicate experiments. The unpaired Student’s t-test was used to determine the significance (* - p ≤ 0.05, ** - p ≤ 0.01, *** - p ≤ 0.001, **** - p ≤ 0.0001) of the differences between the expression profiles of AAV2, UV-AAV2 and scAAVeGFP infected cells (MOI 500). UV-inactivation was assessed on protein level using an anti-Rep antibody (data not shown).

### Cell cycle phase distributions upon AAV2 or rAAV2 infection

Next, the capacity of AAV2 or rAAVeGFP to induce a cell cycle arrest in NHF cells was confirmed by flow cytometry using propidium iodide (PI) staining (Fig. 6, A and C) and confocal laser scanning microscopy (CLSM) using 4‘,6-diamidino-2-phenylindole (DAPI) staining (Fig. 6, B and D). The DAPI based cell cycle analysis workflow was adapted from the protocol published by Roukos et al., 2015 (31) and validated as described previously (32). Both assays revealed an increase in the number of cells in G2 cell cycle phase upon AAV2 (Fig. 6, A and B) or rAAVeGFP (Fig. 6, C and E) infection, indicating a G2 arrest. The shift from S to G2 cell cycle phase upon AAV2 or rAAVeGFP infection was even more pronounced at 48 hpi (see Fig. S3 in the supplemental material).

**FIG 6.**
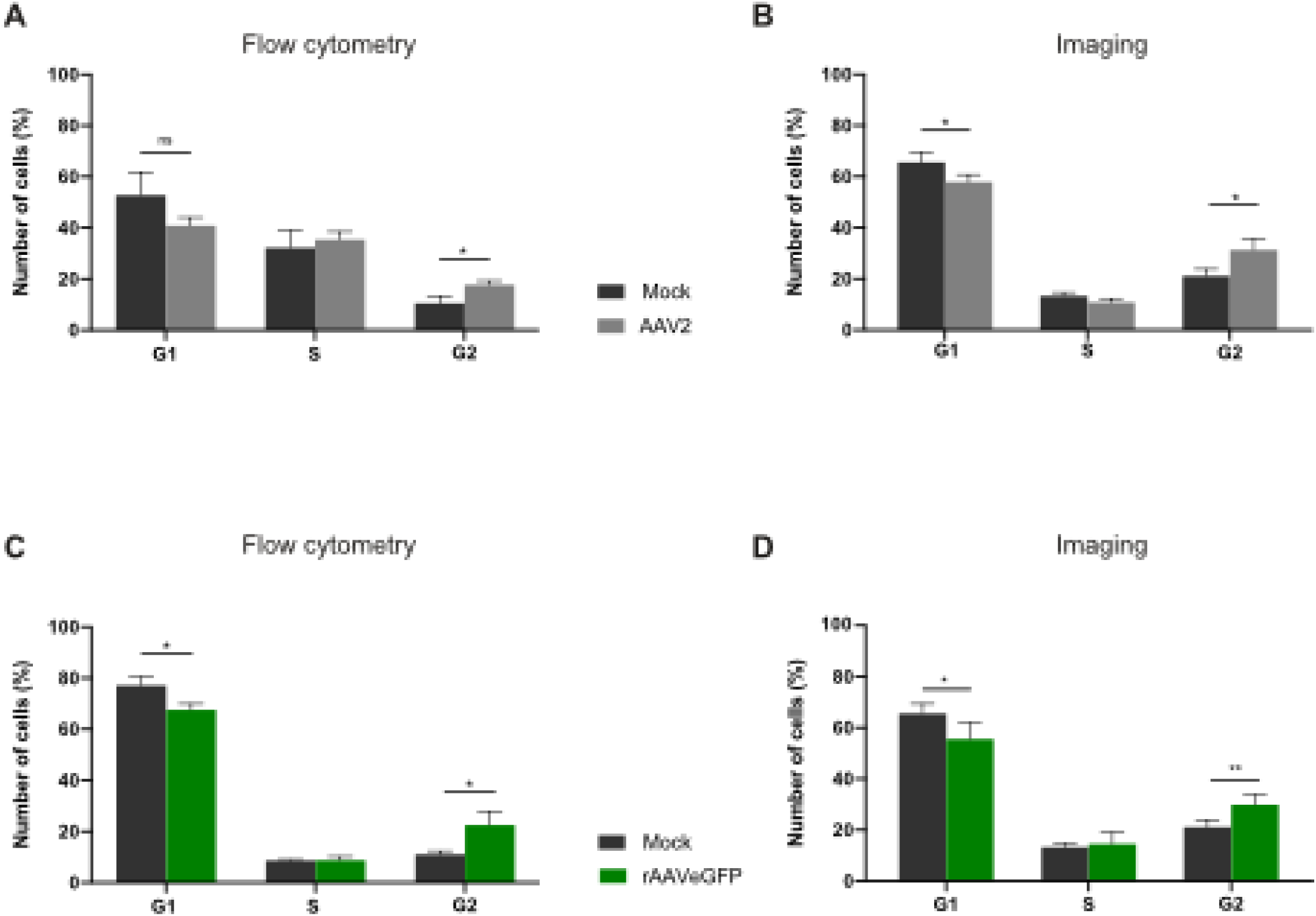
Cell cycle phase distributions upon AAV2 or rAAV2 infection. NHF cells were infected with AAV2 or rAAVeGFP (MOI 5‘000). 24 hpi, the cell cycle profile was assessed either by (A and C) PI staining and flow cytometry (10‘000 cells per sample) or by (B and D) DAPI staining and CLSM (100 cells per sample). Graphs show mean and SD of the percentage of cells in each cell cycle phase. p-values were calculated using an unpaired Student‘s t-test (* - p ≤ 0.05, ** - p ≤ 0.01, *** - p ≤ 0.001, **** - p ≤ 0.0001).

### Upregulated genes in GOterm innate immune response

The upregulated genes in the GOterm innate immune response (Fig. 7) included, among others, the interferon-inducible p200-family protein IFI16, which is assumed to be an innate immune sensor for cytosolic and nuclear ds- as well as ssDNA (13). As the AAV2 DNA can be present as ssDNA, dsDNA and circular dsDNA, AAV2 infection may provoke an IFI16 triggered reaction.

**FIG 7.**
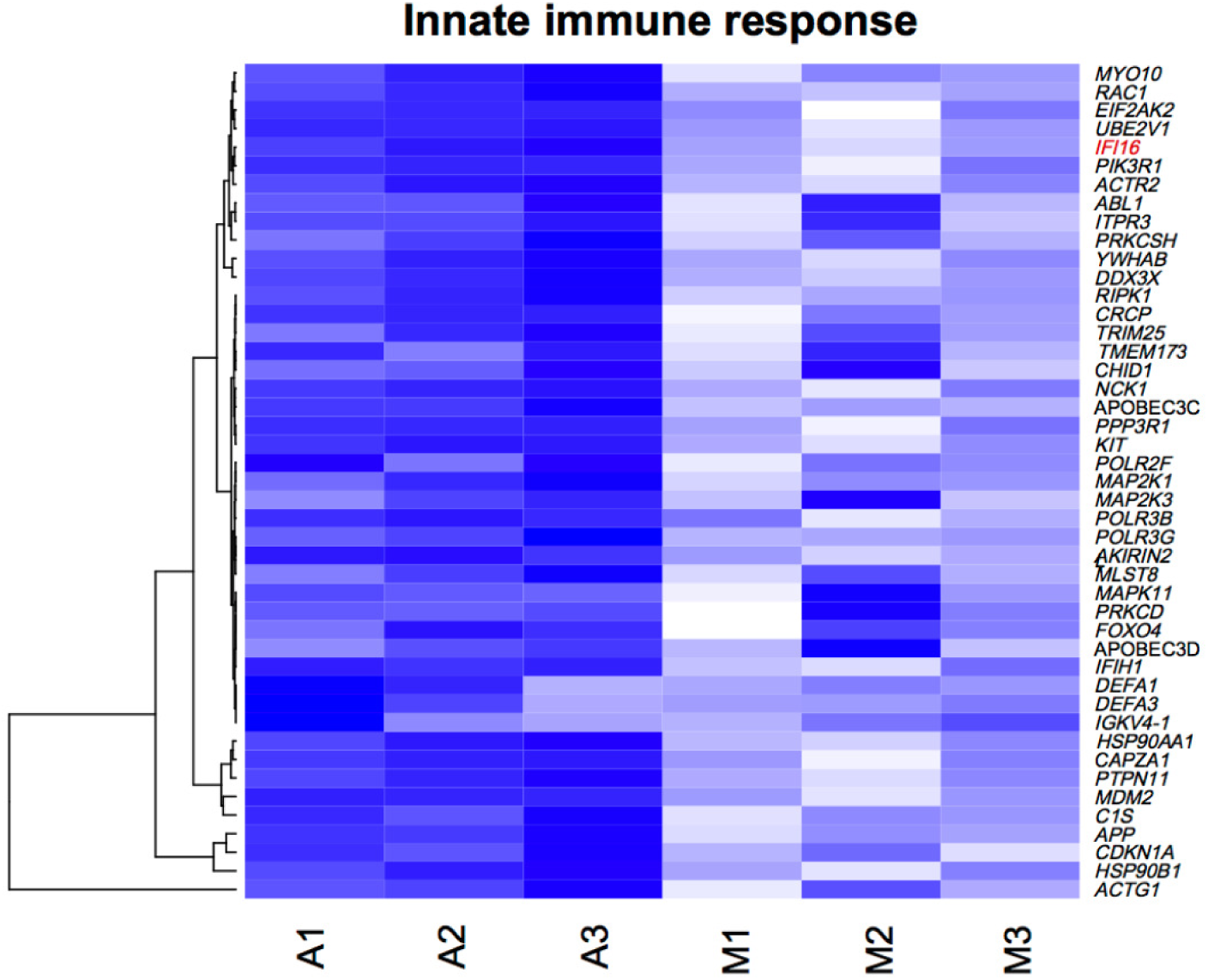
Heat map of upregulated genes in GOterm innate immune response. Reads of the most differentially expressed genes (specified on the right of the heat map) between AAV2-infected (A1-A3) and mock-infected NHF cells (M1-M3). The dendrogram (left side of the heat map) illustrates the unsupervised clustering of the genes. The color key for the log_2_ values of the reads is depicted below. The selected GOI (*IFI16*) is highlighted in red.

### The post-transcriptional silencing of *IFI16* increases AAV2 transduction efficiency independent of the structure of the vector genome

To assess whether IFI16 influences AAV2 vector-mediated cell transduction, NHF cells were transfected with no, scrambled (scr) control or different siRNAs targeting *IFI16*, including a pool of 3 different siRNAs targeting the coding sequence (cds) of *IFI16* as well as a single siRNA targeting the 5’ untranslated region (5‘-UTR) of *IFI16*. At 40 hours post transfection (hpt), cells were either mock-infected or infected with rAAVeGFP or scAAVeGFP, and at 24 hpi, transduced cells counted using a fluorescence microscope (Fig. 8A, B, D, E). Total cell lysates prepared at 24 hpi were subjected to Western blot analysis in order to assess the eGFP (hereinafter referred to as GFP) protein level and to confirm the knock-down of *IFI16* (Fig. 8C and F). C911 siRNA controls were used to confirm that the observed increase of AAV transduction efficiency upon siRNA mediated knock-down of *IFI16* was not due to off-target effects (see Fig. S4 in the supplemental material). One of the C911 siRNA controls, IFI16.3_C911, indeed appeared to knock-down IFI16, at least on protein level (see Fig S4C), suggesting an off-target effect of its corresponding counterpart (IFI16.3_cds) (33). Overall, the data implied an IFI16-mediated inhibition of AAV2 vector-mediated transduction independent of the structure of the vector genome, *i.e.* independent whether a single-stranded or self-complementary vector genome configuration was chosen.

**FIG 8.**
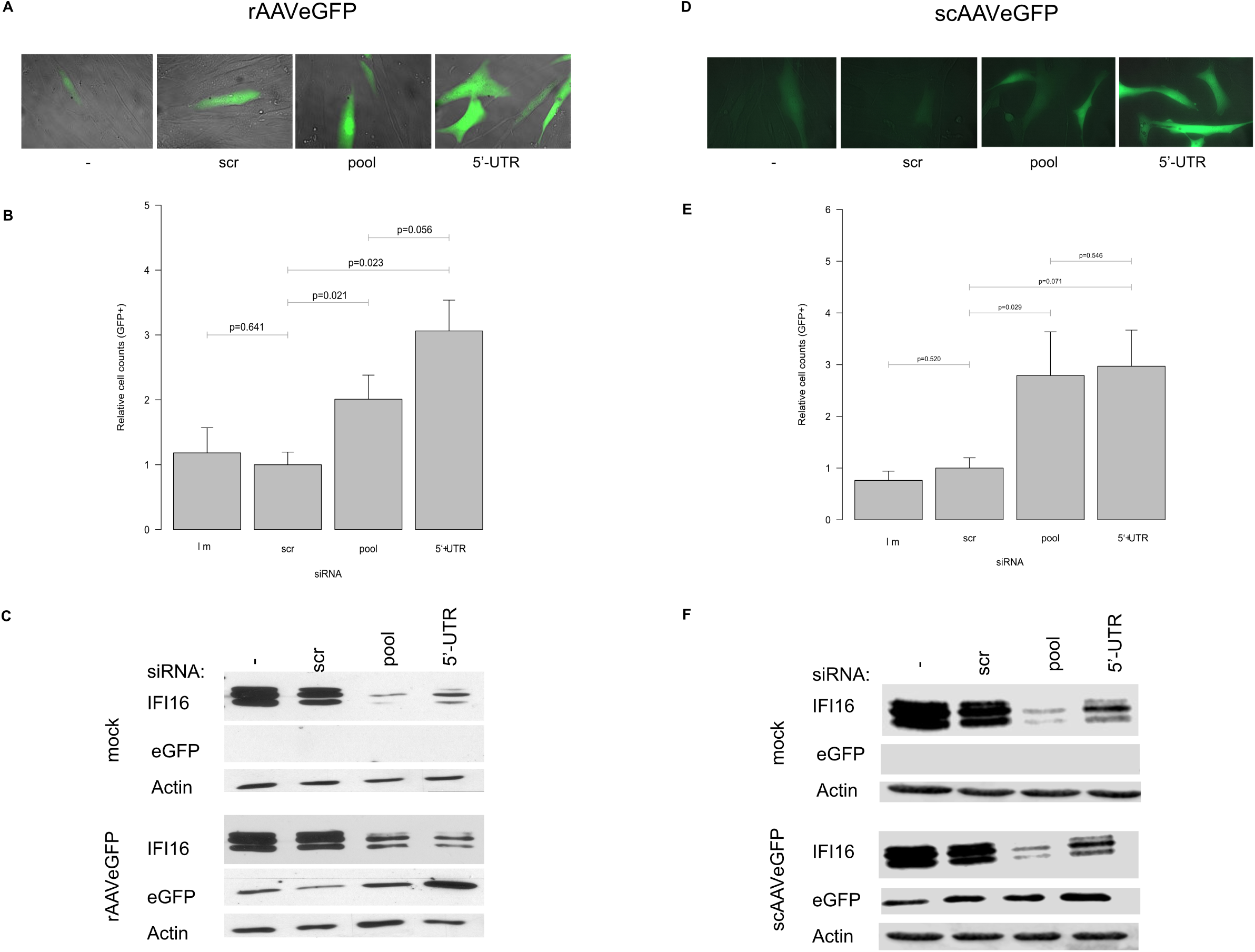
Post-transcriptional silencing of *IFI16* increases AAV2 transduction efficiency independent of the vector genome structure. NHF cells were transfected with no, scr control, or *IFI16* targeting siRNAs. 40 hpt, cells were either mock-infected or infected with rAAVeGFP (MOI 4‘000) or scAAVeGFP (MOI 2‘000). (A and D) 24 hpi, cells were counted using a fluorescence microscope. (B and E) Graphs show mean and SD of the relative cell count of GFP positive NHF cells from triplicate experiments. p-values were calculated using an unpaired Student‘s t-test (* - p ≤ 0.05, ** - p ≤ 0.01, *** - p ≤ 0.001, **** - p ≤ 0.0001). (C and F) Knock-down of *IFI16* was confirmed on protein level (C and F).

### The IFI16-mediated inhibition of AAV2 vector-mediated transduction is interferon signaling independent

To assess whether interferon signaling is relevant for the IFI16-mediated inhibition of AAV2 transduction, 2fTGH Jak1^-/-^ cells were reverse transfected with no siRNA, scr siRNA, a pool of 3 siRNAs targeting the coding sequence of *IFI16* or a single siRNA targeting the 5‘-UTR of *IFI16*. At 40 hpt, the cells were either mock-infected or infected with rAAVeGFP, and at 24 hpi, transduced cells were counted using a fluorescence microscope (Fig. 9A) and subsequently subjected to Western blot analysis to confirm the knock-down of *IFI16* (Fig. 9B). The results in Fig. 9A show an increase in rAAVeGFP transduction efficiency (pool, UTR) also in the Jak1^-/-^ cells, indicating an interferon signaling-independent mechanism of the IFI16-mediated inhibition of AAV2 vector-mediated transduction.

**FIG 9.**
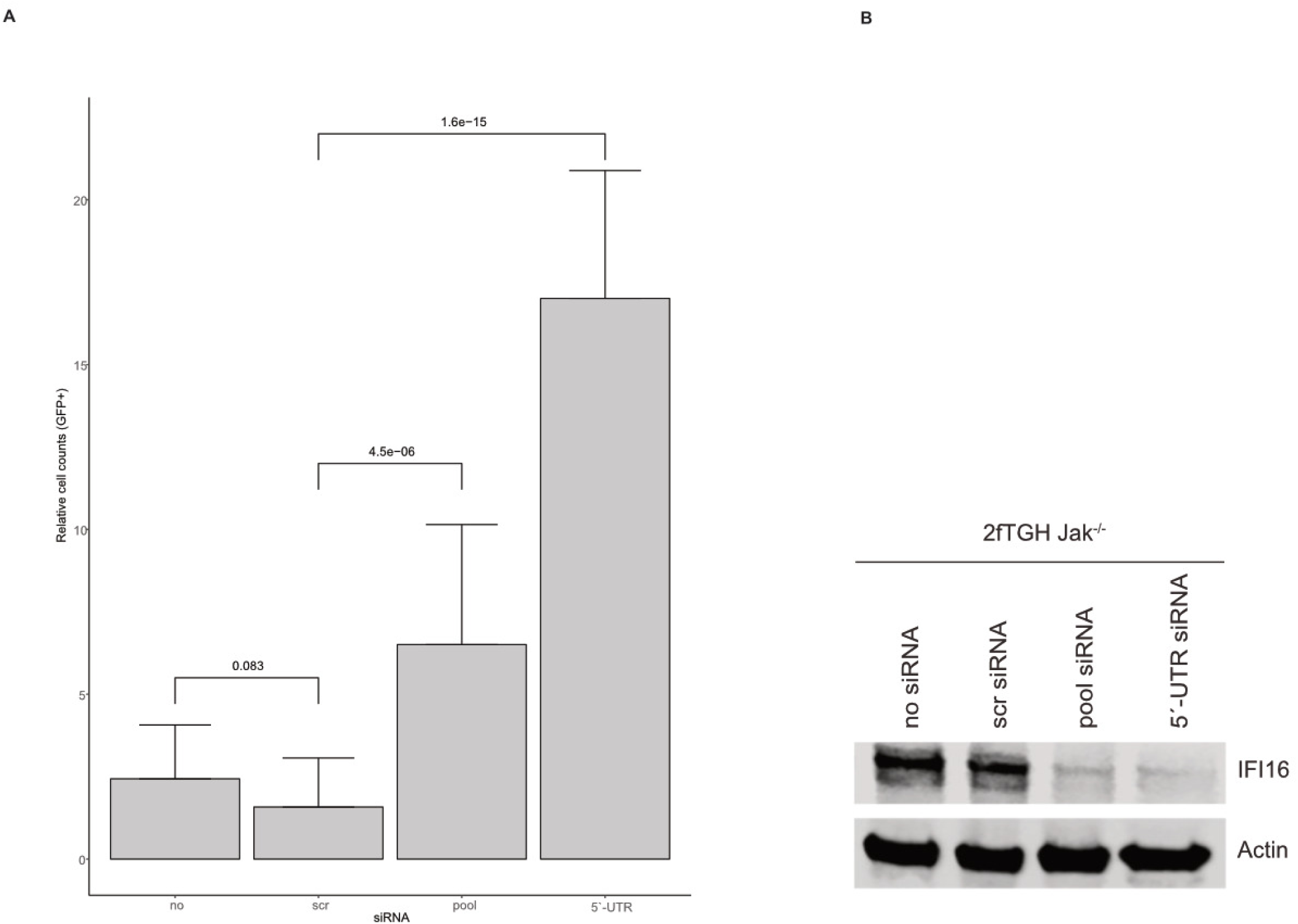
Role of interferon signaling. 2fTGH Jak1^-/-^ cells were transfected with no, scr control, or *IFI16* targeting siRNAs. 40 hpt, cells were either mock-infected or infected with rAAVeGFP (MOI 1‘000). At 24 hpi, transduced cells were counted by fluorescence microscopy. (A) Graph shows mean and SD of the relative cell count of GFP positive 2fTGH Jak1^-/-^ cells from triplicate experiments. p-values were calculated using an unpaired Student‘s t-test (* - p ≤ 0.05, ** - p ≤ 0.01, *** - p ≤ 0.001, **** - p ≤ 0.0001). (B) Knock-down of *IFI16* was confirmed on protein level.

### The IFI16-mediated inhibition of AAV2 vector-mediated transduction is STING independent

The cyclic GMP-AMP synthase (cGAS) - STING (stimulator of interferon genes) pathway plays a crucial role in a variety of viral infections, such as HSV-1, EBV, HPV and HCV (reviewed in (34)). Binding of cGAS to DNA provokes a conformational change of the active site, leading to the synthesis of 2ʹ3ʹ-cyclic GMP-AMP (2ʹ3ʹ-cGAMP). 2ʹ3ʹ-cGAMP operates as a second messenger that binds to the endoplasmic reticulum (ER)-membrane adaptor protein STING and induces a conformational change that results in the activation of STING, leading to the expression of type I interferon and pro-inflammatory cytokines in an IRF-3 or NF-κB dependent manner. Upon induction, IFI16 translocates from the nucleus to the cytoplasm, where it interacts and activates STING (reviewed in (35)). To address the question of functional STING signaling, different cell lines (NHF, U2OS and HeLa) were treated with 2’3’-cGAMP (3 μM) for 9 h, and total RNA was extracted, converted to cDNA and subjected to RT-qPCR using specific primers for *STING* and *ISG56* (see Fig. S5 in the supplemental material). Overall, the results showed no impairment of the STING pathway in NHF cells, whereas in U2OS and HeLa cells a deficient STING pathway was observed. To explore the role of STING in the IFI16-mediated inhibition of AAV vector transduction, NHF cells were transfected with no, scr or different siRNAs, including a pool of 3 different siRNAs targeting *IFI16*, as well as single siRNAs targeting the cds of *STING*. At 40 hpt, the cells were either mock-infected or infected with rAAVeGFP or scAAVeGFP, and at 24 hpi, the transduced cells were counted by using a fluorescence microscope (Fig. 10A) and subsequently subjected to Western blot analysis to confirm the knock-down of *IFI16* and *STING* (Fig. 10B). Generally, the data indicated a STING-independent IFI16-mediated inhibition of AAV2 vector-mediated transduction.

**FIG 10.**
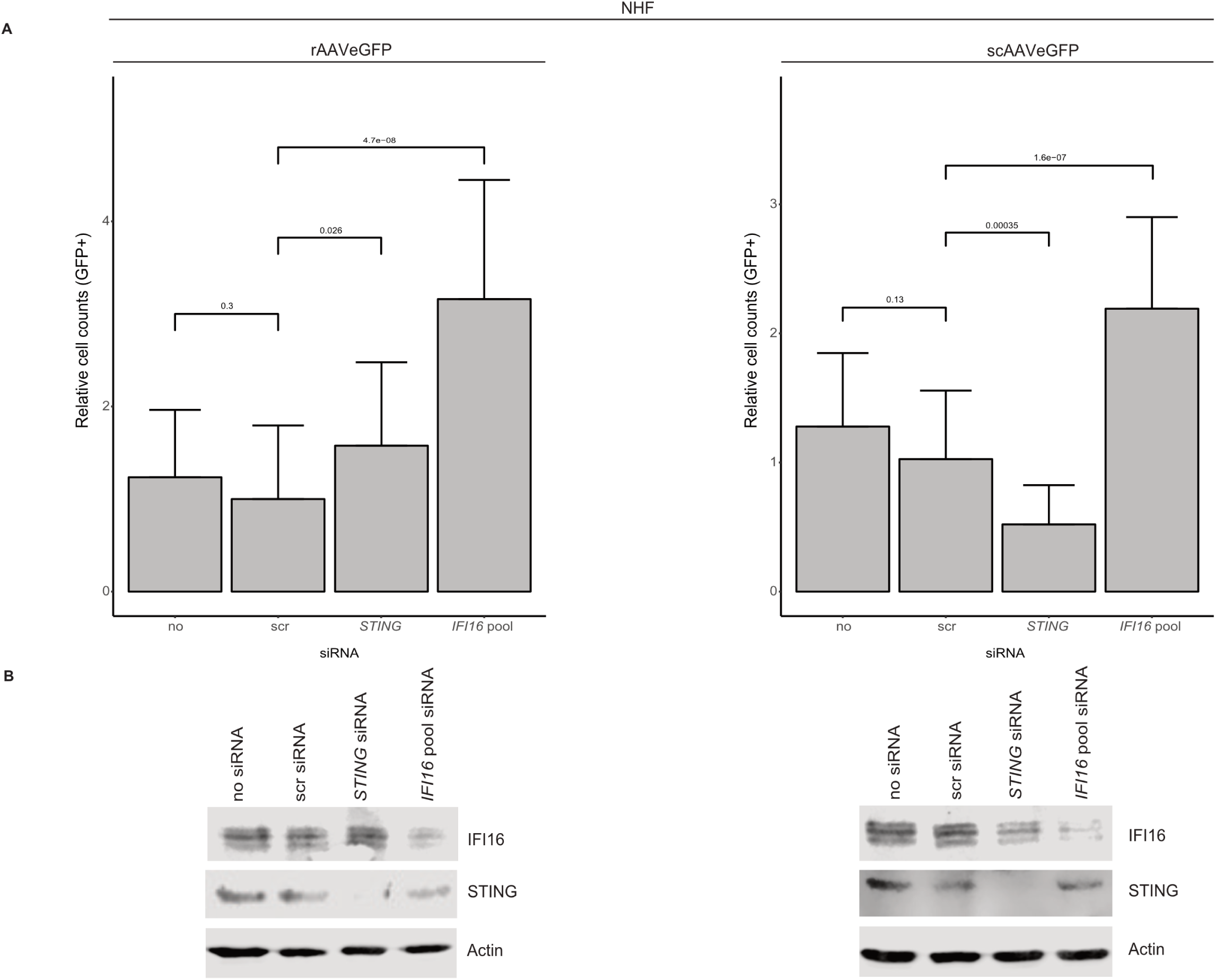
Effect of STING on AAV2 vector-mediated transduction. NHF cells were transfected with no, scr control, *STING* or *IFI16* targeting siRNAs. 40 hpt, cells were either mock-infected, infected with rAAVeGFP (MOI 6‘000) or scAAVeGFP (MOI 4‘000). 24 hpi, transduced cells were counted by fluorescence microscopy. (A) Graphs show mean and SD of the relative cell count of GFP positive NHF cells from triplicate experiments. p-values were calculated using an unpaired Student‘s t-test (* - p ≤ 0.05, ** - p ≤ 0.01, *** - p ≤ 0.001, **** - p ≤ 0.0001). (B) Knock-down of *IFI16* and *STING* was confirmed on protein level.

### Exogenous complementation of *IFI16* in U2OS *IFI16*^-/-^ cells

With respect to the impaired STING signaling in U2OS cells, an assay was established to exogenously complement *IFI16* in U2OS *IFI16*^-/-^ cells. To this end, U2OS IFI16^-/-^ cells were either untransduced (no), transduced with lentiviral vectors expressing GFP (GFP ctrl.), or transduced with lentiviral vectors expressing *IFI16* fused to monomeric GFP (IFI16_GFP). 72 hours later, the cells were infected with rAAVmCherry and, 24 hpi, the cells were subjected to RT-qPCR (Fig. 11A). Overall, the results showed a decrease in the relative mCherry expression (Fig. 11A) upon exogenous complementation of *IFI16* (Fig. 11B).

**FIG 11.**
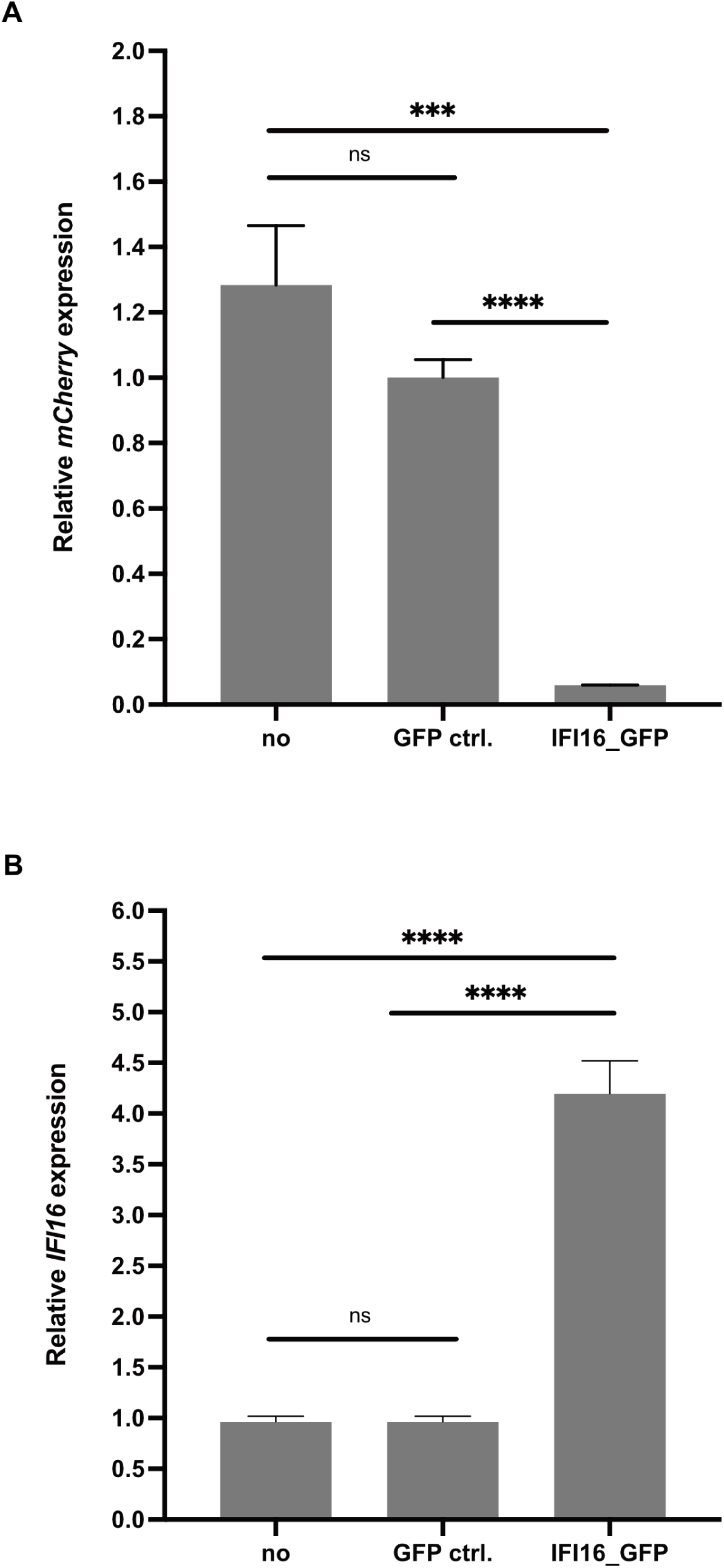
Exogenous complementation of *IFI16* in U2OS *IFI16*^-/-^ cells. U2OS IFI16^-/-^ were either untransduced, transduced with lentiviral vectors expressing GFP (MOI 5), or transduced with lentiviral vectors expressing *IFI16* fused to monomeric GFP (MOI 5) in presence of polybrene. 72 hours later, the cells were infected with rAAV2mCherry (MOI 500). (A) mCherry expression was assessed by RT-qPCR using specific primers for mCherry. (B) Exogenous complementation of *IFI16* was confirmed on transcript level. Graphs show mean and SD of the relative gene expression from triplicate experiments. p-values were calculated using an unpaired Student‘s t-test (* - p ≤ 0.05, ** - p ≤ 0.01, *** - p ≤ 0.001, **** - p ≤ 0.0001).

### Sub-nucleolar localization of IFI16

To assess the spatial distribution of IFI16 and AAV2 genomes a combined immunofluorescence analysis (IF) and fluorescence *in situ* hybridization (FISH) was established. To this end, NHF cells were infected with AAV2 and 24 hpi fixed and processed for multicolor IF analysis combined with FISH and CLSM. The results showed the accumulation of IFI16 in nucleoli, together with AAV2 DNA and, conditionally, AAV2 capsids (Fig. 12, A-C; and Fig. S6 in the supplemental material). However, the image-based quantification of the post-transcriptional silencing of *IFI16* revealed that neither partial uncoating in the cytoplasm, complete uncoating in the nucleolus, or cell cycle progression (32) was affected (data not shown).

**FIG 12.**
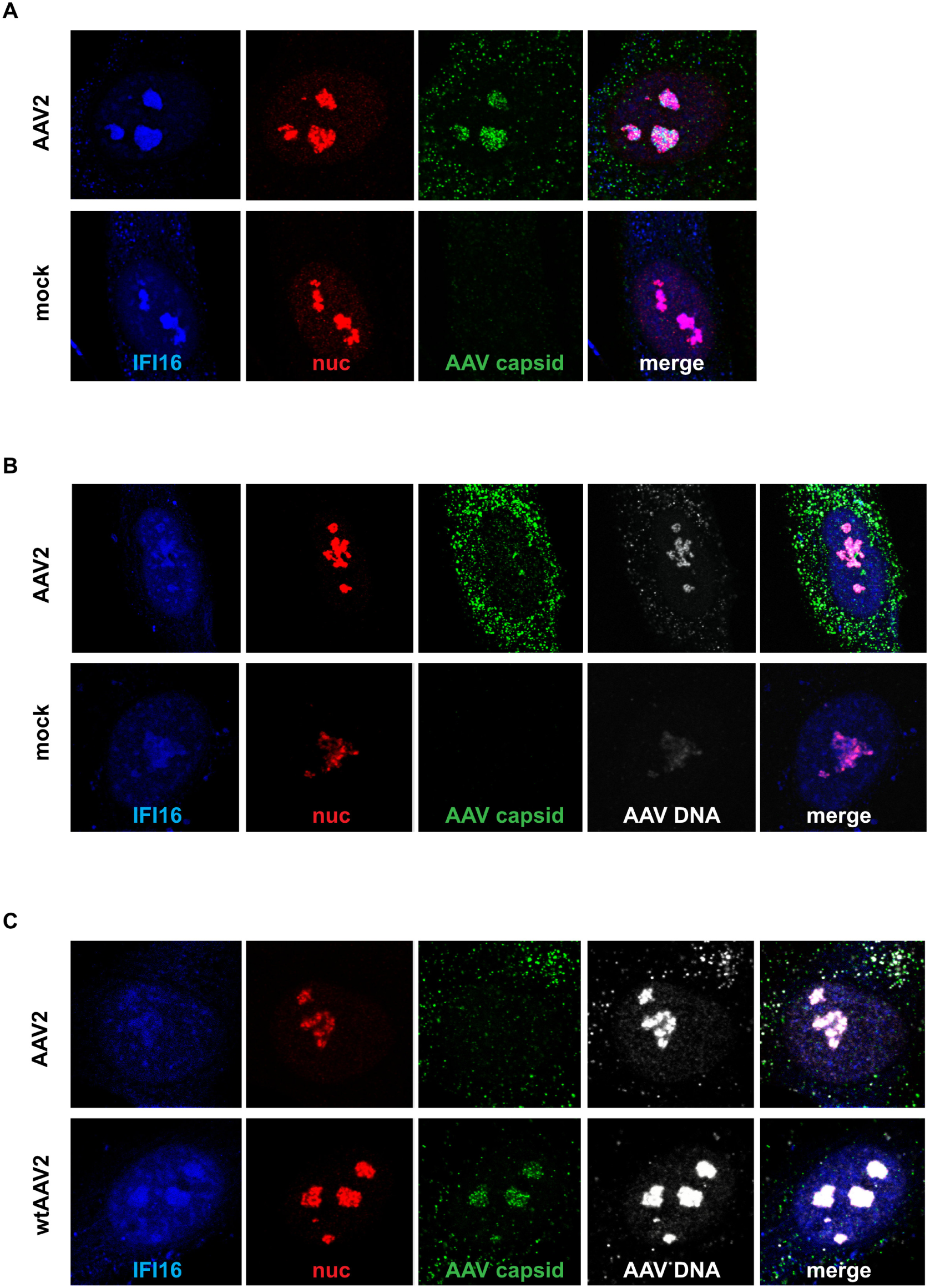
Multicolor IF combined with FISH. NHF cells were infected with AAV2 (MOI 20‘000). After 24 h, the cells were fixed and processed for multicolor IF analysis combined with FISH and CLSM. IFI16 was detected by direct labeling of the antibody to ATTO-390 (blue). Nucleoli were visualized using an antibody against fibrillarin (red). Capsids were detected using an antibody against intact AAV2 capsids (green). AAV2 DNA (grey) was detected by linking the amine-modified DNA to AF647. (A) Nucleolar localization of IFI16 and AAV2 capsids. (B) Nucleolar localization of IFI16 and AAV2 DNA. (C) Nucleolar localization of IFI16, AAV2 DNA and, conditionally, intact AAV2 capsids.

### The post-transcriptional silencing of *IFI16* increases AAV2 *rep* but not *cap* expression

To explore the influence of IFI16 on the expression of AAV2 specific genes, *rep* and *cap*, NHF (Fig. 13A) and U2OS (Fig. 13B) cells were reverse transfected with either scr siRNA or different siRNAs, including a pool of 3 different siRNAs targeting cds of *IFI16* (pool), as well as single siRNA targeting the 5‘-UTR of *IFI16*. At 40 hpt, cells were infected with AAV2 and 24 hours later, total RNA was extracted and subjected to RT-qPCR using specific primers for the Rep helicase domain (*rep*), *cap* gene (*cap*), or *IFI16*. In summary, the data showed an increase in *rep* expression upon knock-down of *IFI16* in both NHF and U2OS cells, indicating a putative interplay of IFI16 and AAV2 gene expression. However, *cap* expression was neither affected in NHF nor in U2OS cells upon the post-transcriptional silencing of *IFI16*. Besides, the post-transcriptional silencing of *IFI16* did not only result in an increase of *rep* expression but also enhanced AAV2 genome replication in presence of adenovirus type 5 (AdV5; Fig. 14C) without affecting the relative genome copy numbers of AAV2 (Fig. 14A) or AdV5 (Fig. 14B).

**FIG 13.**
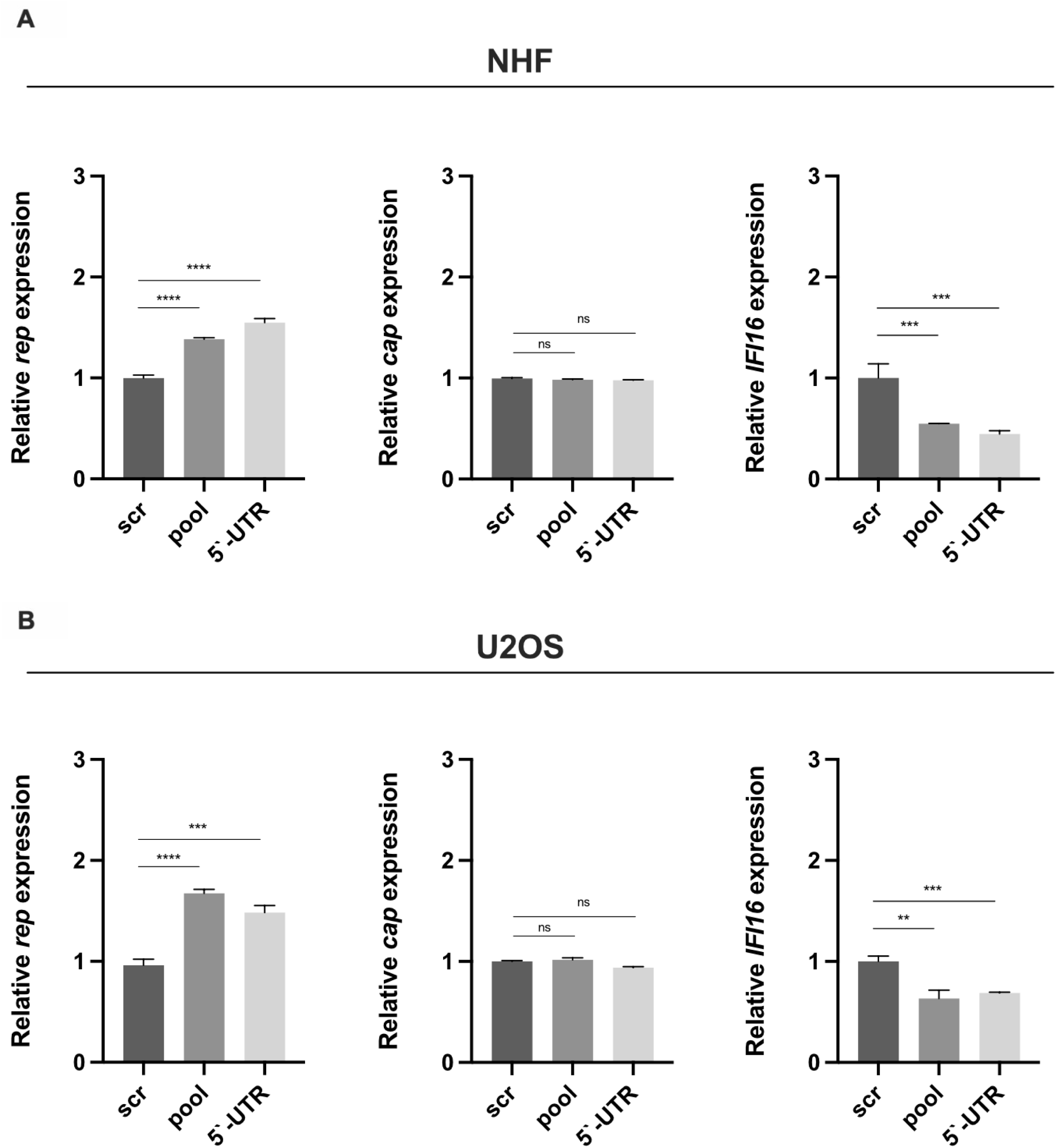
Post-transcriptional silencing of *IFI16* increases AAV2 *rep* but not *cap* expression. (A) NHF and (B) U2OS cells were transfected with scr control or *IFI16* targeting siRNAs, respectively. 40 hpt, cells were infected with AAV2 (NHF; MOI 4‘000, U2OS; MOI 2‘000). 24 hpi, total RNA was extracted and subjected to RT-qPCR using specific primers for the Rep helicase domain (*rep*), *cap* gene (*cap*), or *IFI16*. Graphs show mean and SD of the relative gene expression from triplicate experiments. p-values were calculated using an unpaired Student‘s t-test (* - p ≤ 0.05, ** - p ≤ 0.01, *** - p ≤ 0.001, **** - p ≤ 0.0001).

**FIG 14.**
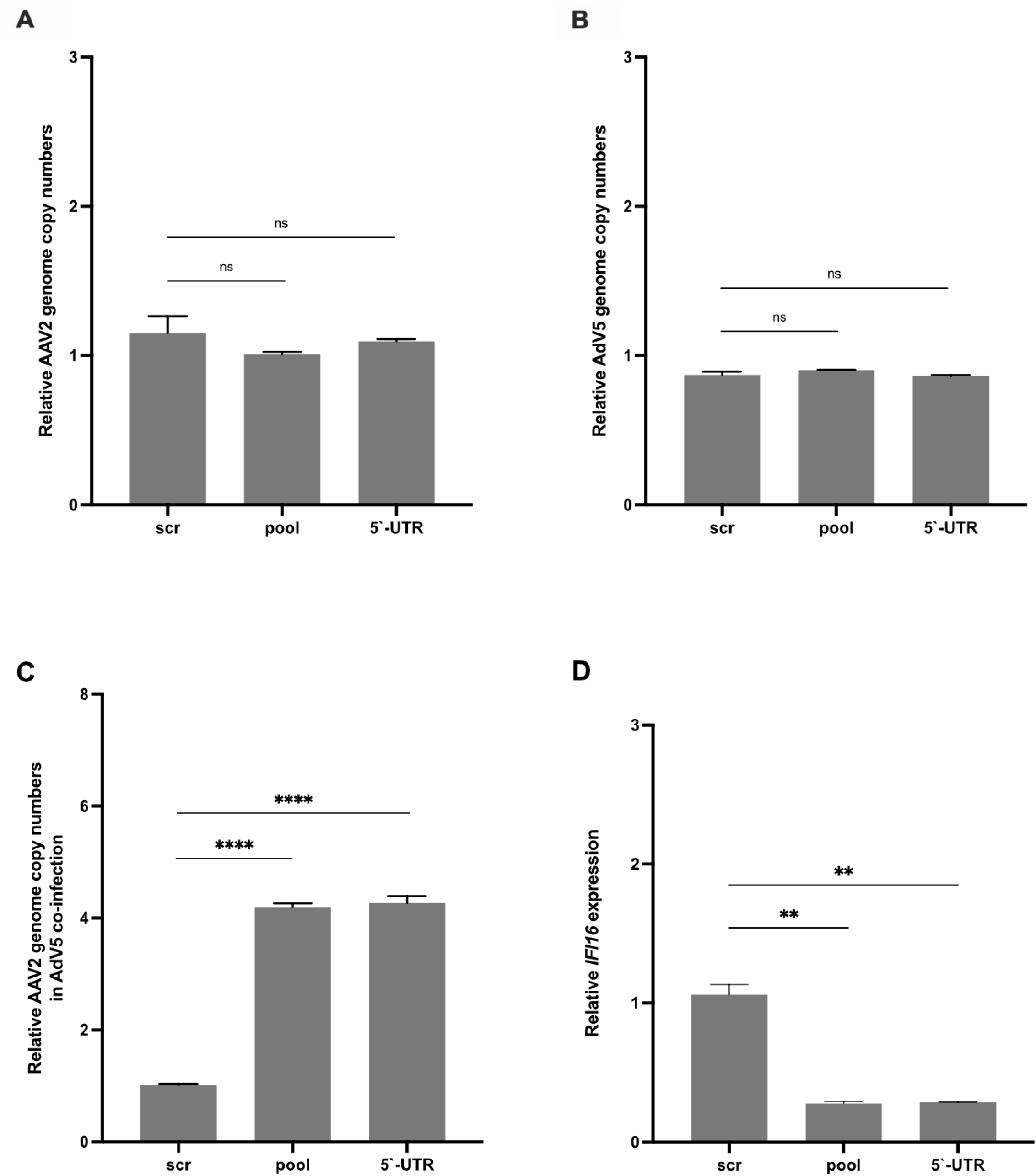
Post-transcriptional silencing of *IFI16* increases AAV2 genome replication in presence of AdV5. NHF cells were transfected with scr control or *IFI16* targeting siRNAs, respectively. 40 hpt, cells were either infected with (A) AAV2 (MOI 2‘000), (B) AdV5 (MOI 5), or (C) co-infected with AAV2 (MOI 2‘000) and AdV5 (MOI 5). After 24 h, total DNA was isolated and subjected to qPCR using specific primers for AAV2 or AdV5, respectively. (D) Knock-down of *IFI16* was confirmed on transcript level. Graphs show mean and SD of the relative genome copy numbers or the relative gene expression, respectively, from triplicate experiments. p-values were calculated using an unpaired Student‘s t-test (* - p ≤ 0.05, ** - p ≤ 0.01, *** - p ≤ 0.001, **** - p ≤ 0.0001).

### The post-transcriptional silencing of *IFI16* increases vector-mediated GFP expression

To assess the influence of IFI16 on vector-mediated GFP expression, NHF and U2OS cells were reverse transfected with either scr siRNA, or different siRNAs, including a pool of 3 different siRNAs targeting the cds of *IFI16* (pool), as well as a single siRNA targeting the 5‘-UTR of *IFI16*, or a control siRNA targeting GFP. At 40 hpt, cells were either infected with rAAVeGFP (Fig. 15) or scAAVeGFP (Fig. 16) and 24 hours later, total RNA was extracted and subjected to RT-qPCR using specific primers for GFP or *IFI16*. In summary, the data showed a strong increase in GFP expression upon knock-down of *IFI16* in both cell lines regardless of the structure of the vector genome, single-stranded or self-complementary genome configuration.

**FIG 15.**
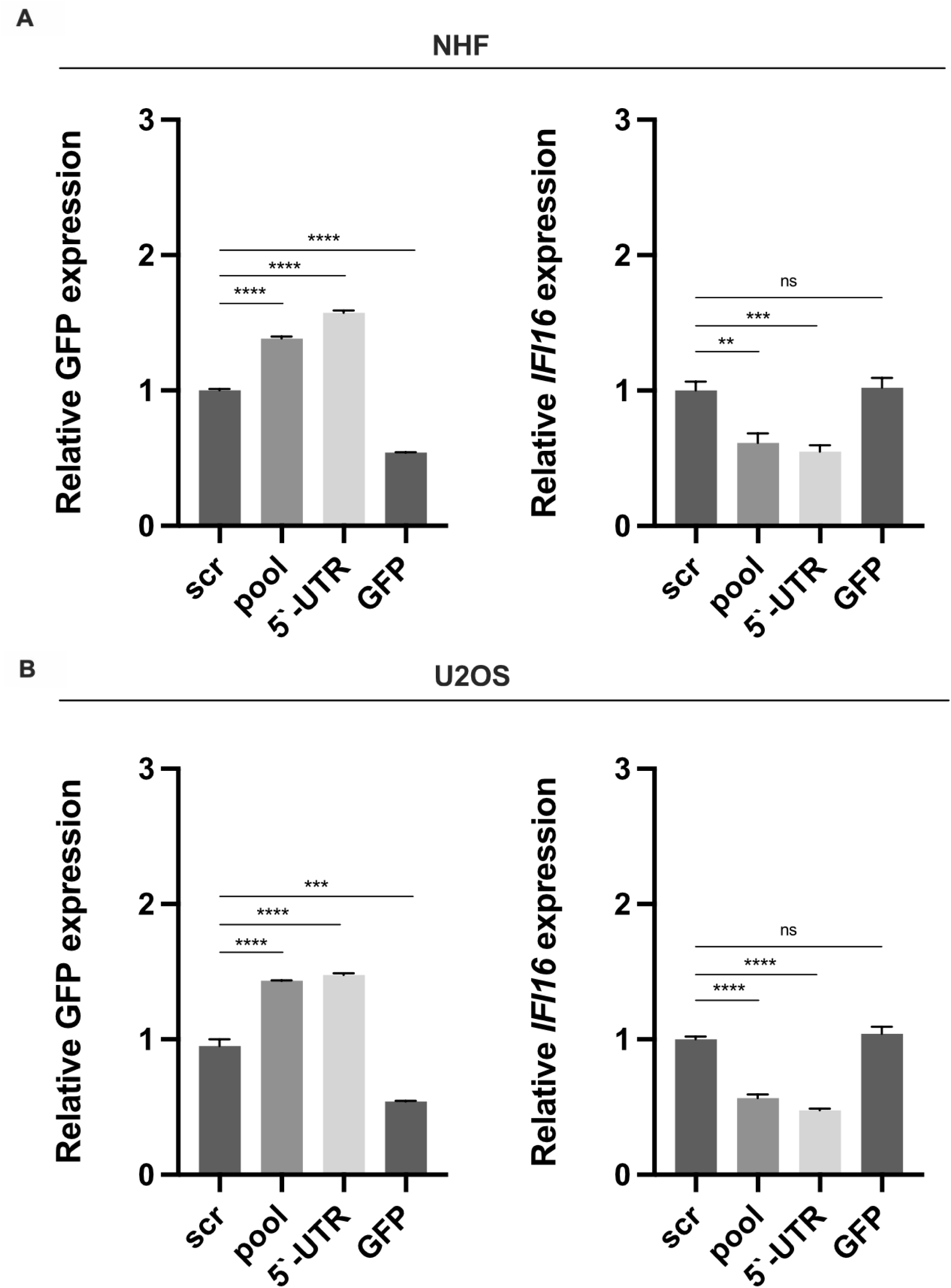
Post-transcriptional silencing of *IFI16* increases single-stranded rAAV2 vector-mediated GFP expression. (A) NHF and (B) U2OS cells were transfected with scr control, GFP control, or *IFI16* targeting siRNAs. 40 hpt, cells were infected with rAAVeGFP (NHF; MOI 4‘000, U2OS; MOI 2‘000). 24 hpi, total RNA was extracted and subjected to RT-qPCR using specific primers for GFP or *IFI16*. Graphs show mean and SD of the relative gene expression from triplicate experiments. p-values were calculated using an unpaired Student‘s t-test (* - p ≤ 0.05, ** - p ≤ 0.01, *** - p ≤ 0.001, **** - p ≤ 0.0001).

**FIG 16.**
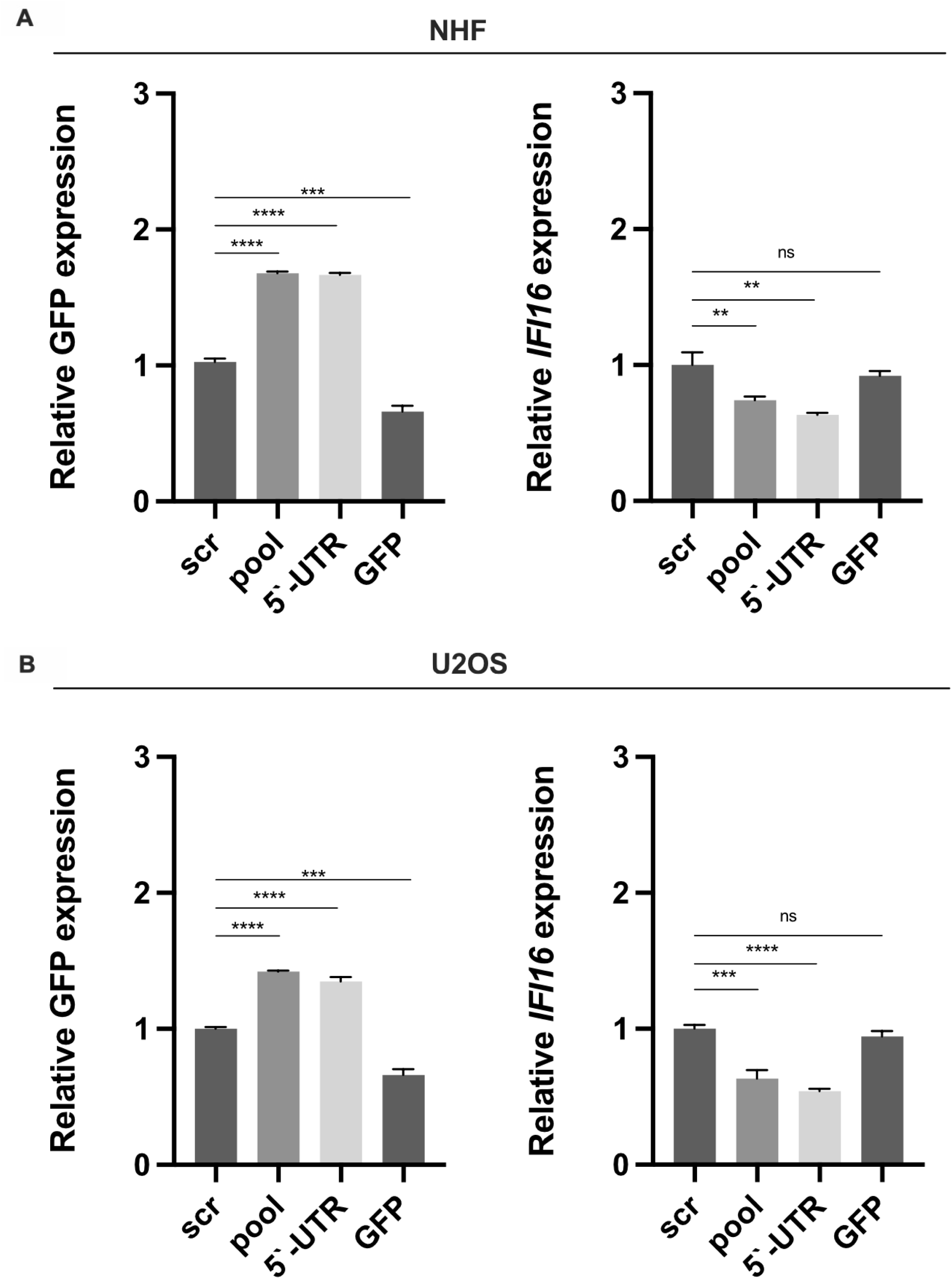
Post-transcriptional silencing of *IFI16* increases self-complementary scAAV2 vector-mediated GFP expression. A) NHF and (B) U2OS cells were transfected with scr control, GFP control, or *IFI16* targeting siRNAs. 40 hpt, cells were infected with rAAVeGFP (NHF; MOI 2‘000, U2OS; MOI 1‘000). 24 hpi, total RNA was extracted and subjected to RT-qPCR using specific primers for GFP or *IFI16*. Graphs show mean and SD of the relative gene expression from triplicate experiments. p-values were calculated using an unpaired Student‘s t-test (* - p ≤ 0.05, ** - p ≤ 0.01, *** - p ≤ 0.001, **** - p ≤ 0.0001).

### IFI16 inhibits AAV2 gene expression in an Sp1-dependent manner

Emerging evidence imply that IFI16 exerts its effects via various genome regulation mechanisms, independently of innate immune sensing. For example, IFI16 promotes the addition of heterochromatin marks and yet reduces the number of euchromatin marks on specific viral genomes (15, 16) or it affects viral gene expression by reducing the availability of the transcription factor Sp1 (19).

As both wild-type AAV2 and AAV2 vector genome expression was affected by the post-transcriptional silencing of *IFI16,* and as we did not observe any changes in methylation marks on either the wild-type or the vector genome (data not shown), we assessed whether IFI16 exerts its effect in a Sp1-dependent manner. To this end, U2OS IFI16^-/-^ or U2OS wt were either untransduced (U2OS IFI16^-/-^ or U2OS wt, respectively), transduced with lentiviral vectors expressing GFP (U2OS IFI16^-/-^ + GFP ctrl.), or transduced with lentiviral vectors expressing *IFI16* fused to monomeric GFP (U2OS IFI16^-/-^ + IFI16_GFP) in presence of polybrene. 72 hours later, the cells were either mock-infected or infected with AAV2 (MOI 20‘000) and 24 h later subjected to chromatin immunoprecipitation (ChIP) assays using an anti-Sp1 antibody and primers for the p5 (Fig. 17A) or p19 (Fig. 17B) promoter regions, respectively. Intriguingly, the relative Sp1 promoter occupancy of p5 and p19 was significantly higher in the untransduced (U2OS IFI16^-/-^) or control vector transduced cells (U2OS IFI16^-/-^ + GFP ctrl.) compared to the *IFI16* complemented cells (U2OS IFI16^-/-^ + IFI16_GFP) or the parental cell line (U2OS wt), respectively, indicating that IFI16 indeed restricts AAV2 independently of immune sensing by binding (Fig. 17C) and inhibiting the host transcription factor Sp1 that transactivates the viral promoter regions, thereby driving AAV2 gene expression. Moreover, the IFI16-mediated unavailability of Sp1 did not only affect the AAV2 rep promoters but also the CMV promoter in AAV2 vector genomes (Fig. 18). However, the relative CMV promoter occupancy of the self-complementary AAV2 vector (Fig. 18B) was less pronounced compared to the single-stranded vector (Fig. 18A), which might be reasoned by the nature of this promoter (minimal-CMV), possessing less Sp1 binding sites than the full length CMV promoter of the single-stranded AAV2 vector. Overall, this data implies that IFI16 inhibits gene expression of wild-type AAV2 and AAV2 vectors by reducing the availability of the transcription factor Sp1, thereby reducing the SP1-mediated transactivation of the viral promoters.

**FIG 17.**
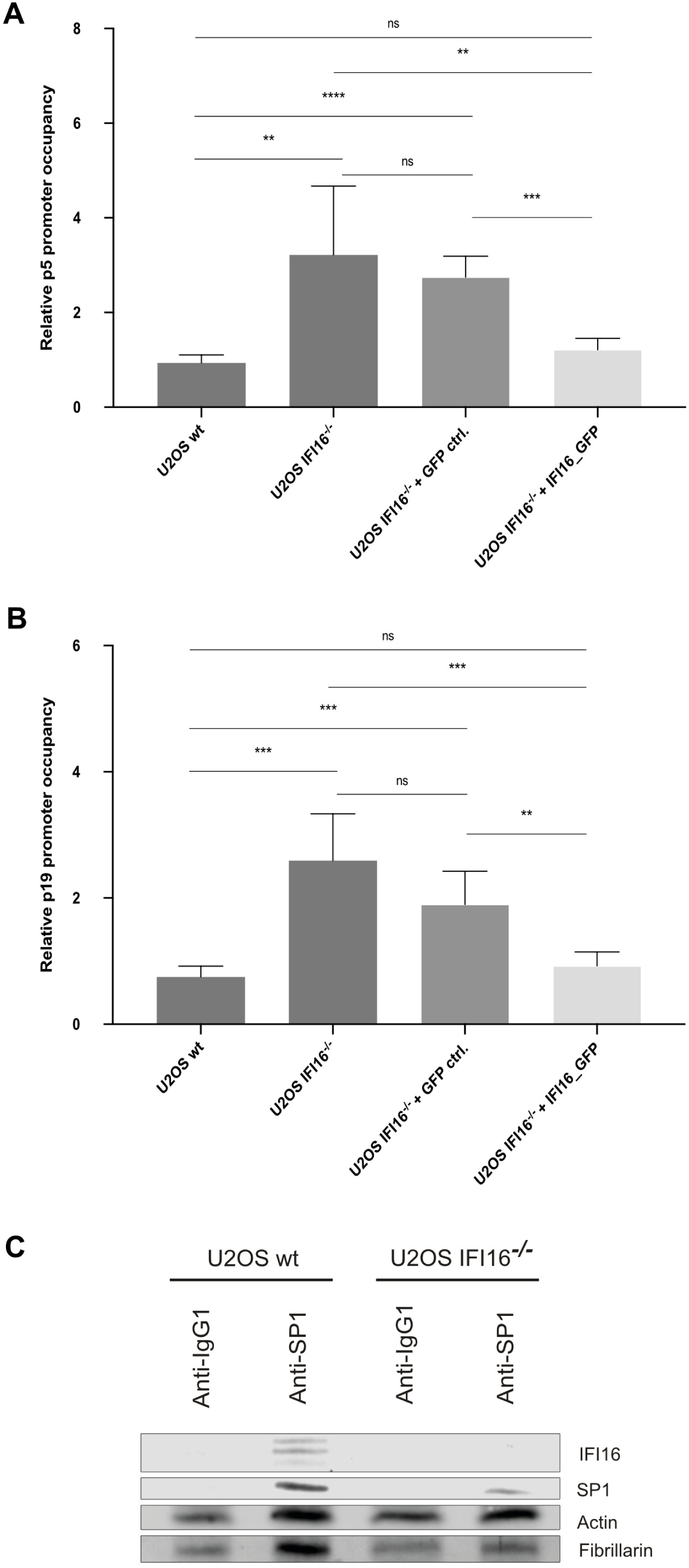
IFI16 inhibits AAV2 gene expression in an Sp1-dependent manner. U2OS IFI16^-/-^ and U2OS wt cells were either untransduced (U2OS IFI16^-/-^ or U2OS wt, respectively), transduced with lentiviral vectors expressing GFP (MOI 5; U2OS IFI16^-/-^ + GFP ctrl.), or transduced with lentiviral vectors expressing *IFI16* fused to monomeric GFP (MOI 5; U2OS IFI16^-/-^ + IFI16_GFP). 72 hours later, the cells were infected with AAV2 (MOI 20‘000) and 24 h later subjected to ChIP assays using an anti-Sp1 antibody and primers for the (A) p5 or (B) p19 promoter regions, respectively. (C) Interaction of IFI16 and Sp1 was assessed by Co-IP in U2OS wt and U2OS IFI16^-/-^ cells. Graphs show mean and SD of the relative promoter occupancy from triplicate experiments. p-values were calculated using an unpaired Student‘s t-test (* - p ≤ 0.05, ** - p ≤ 0.01, *** - p ≤ 0.001, **** - p ≤ 0.0001).

**FIG 18.**
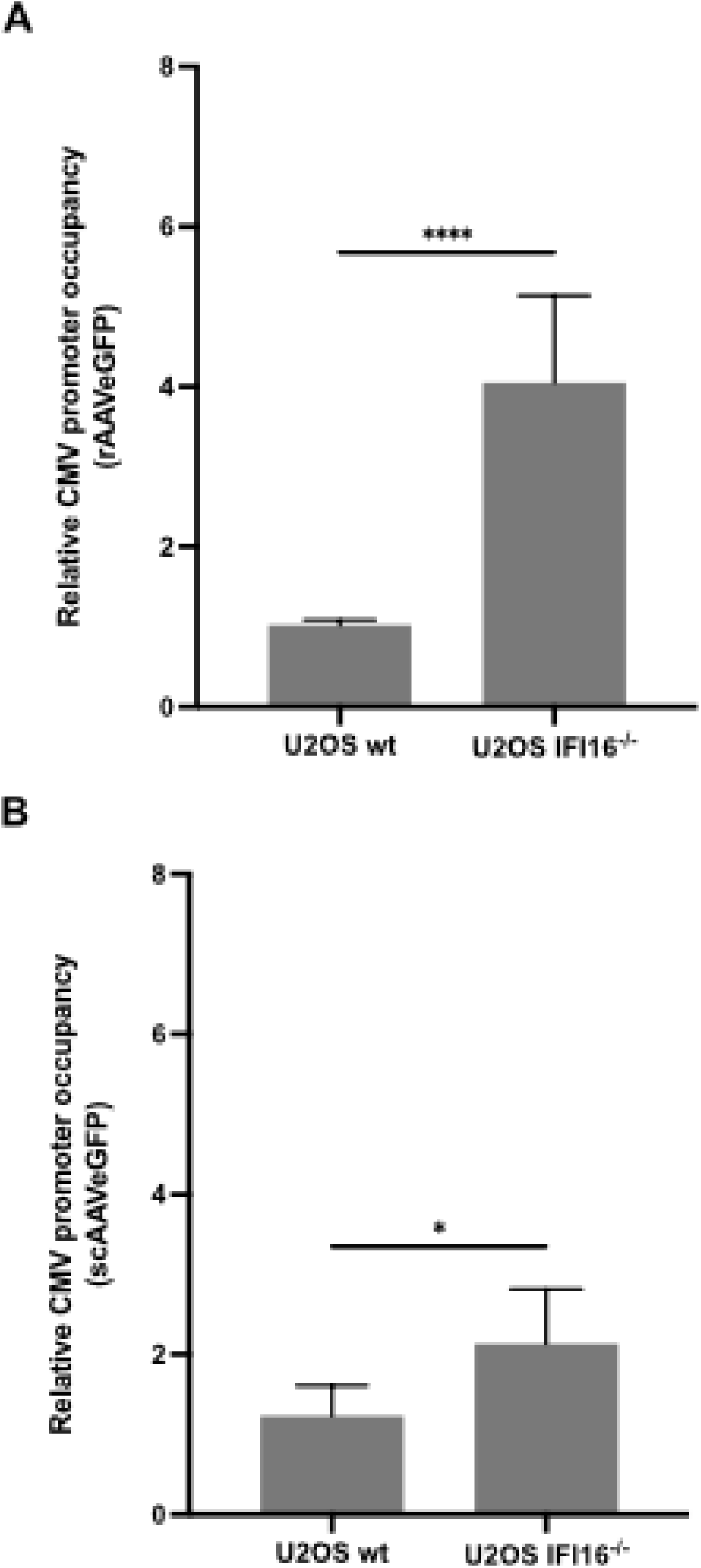
IFI16 inhibits AAV2 vector-mediated gene expression in an Sp1-dependent manner. U2OS IFI16^-/-^ cells or the parental cell line U2OS wt were infected with (A) rAAVeGFP (MOI 20‘000) or (B) scAAVeGFP (MOI 20‘000) and, 24 hpi, the cells were subjected to ChIP assays using an anti-Sp1 antibody and primers for CMV promoter regions. Graph shows mean and SD of the relative promoter occupancy from triplicate experiments. p-values were calculated using an unpaired Student‘s t-test (* - p ≤ 0.05, ** - p ≤ 0.01, *** - p ≤ 0.001, **** - p ≤ 0.0001).

## DISCUSSION

Downstream analysis of the RNA-seq data that included 1‘930 genes with a p-value < 0.01 and more than 40 reads showed 8 distinct clusters of biological processes, which differ between AAV2 and mock-infected NHF cells. The most prominent biological processes include the regulation of macromolecules / metabolic processes / gene expression and cell cycle regulation. Further analysis of the clusters with the aid of heat maps resulted in the identification of the 50 most differentially expressed genes. Of special note was the heat map for the chromatin organization, as it showed a maximum of difference in reads (log_2_) of roughly |2|, representing a fold difference of 4 between AAV2 and mock-infected cells. A possible explanation for the down regulation of many histones could be that the histone deacetylase-2 (*HDAC2*) and the NAD-dependent deacetylase sirtuin-1 (*SIRT1*) genes are up-regulated. Histone deacetylases remove acetyl groups, which results in an increase of positively charged histone tails. This leads to a high-affinity binding between the histones and DNA backbone, entailing in a condensed DNA structure, which prevents transcription. Further evidence was given by several up-regulated histone methyltransferases, which transfer methyl groups from S-adenosyl methionine (SAM) either on arginine or lysine residues of the H3 and H4 histones. Methylated histones can then either be transcriptionally active or repressed. It would be interesting to further evaluate the data in order to draw a conclusion whether the found methyltransferases act as silencers or activators and whether or not this aforesaid AAV2-induced histone modification profiles are due to the AAV2-mediated cell cycle arrest.

When excluding the genes with a fold change of < |1.5|, the list with the 1’930 DE genes was reduced to 872 DE genes, of which 268 were down-regulated and 604 up-regulated. To gain a first impression in terms of biological relevance, the 872 genes were projected onto KEGG pathways. The KEGG analysis revealed that around 80% of the genes of the surveillance system of the cell cycle are negatively regulated upon AAV2 infection. To further evaluate the 872 differentially expressed genes, 10 genes were selected due to their differential expression profile in AAV2-infected and mock-infected cells and based on their relevance in the before mentioned downstream analysis. Both, transcription and protein levels of the selected genes assessed by RT-qPCR and Western blot analysis, respectively, correlated well with the RNA-seq and connoted a differential gene regulation upon AAV2 infection.

Moreover, the expression profiles of wild-type AAV2 and UV-inactivated AAV2-infected cells indicated a shift in cell cycle progression upon infection, while that of cells infected with a recombinant AAV2 with a self-complementary genome configuration (scAAVeGFP) did not. This finding is in accordance with previous observations (30), showing that infection with self-complementary AAV2 vectors allowed the cells to progress through mitosis, an event that occurred significantly less frequently upon infection with single-stranded recombinant AAV2 (rAAV2) vectors. However, recombinant single-stranded AAV2 vectors, similar to wild-type AAV2 (36), may induce cell cycle arrest more efficiently compared to self-complementary AAV2 vectors due to the single-stranded nature of the genome.

Further examination of the differentially expressed genes in the different GOterms revealed an upregulation of several genes in the GOterm innate immune response, including the interferon-inducible p200-family protein IFI16, which is assumed to be an innate immune sensor for cytosolic and nuclear ds- as well as ssDNA (13). IFI16 has been shown to be a restriction factor of many different viruses through several mechanisms, including epigenetic modifications and interferon response. For example, it was noted that IFI16 acts as a cytosolic immune sensor of HIV-1 DNA species in macrophages (37) and promotes interferon induction via the cyclic GMP-AMP synthase (cGAS) and stimulator of interferon genes (STING) pathway (38). Another study reported the IFI16-mediated sensing of HIV-1 reverse transcription intermediates, resulting in a caspase-1-dependent pyroptotic cell death of HIV infected CD4+ T cells (39).

Although IFI16 is thought to act as a cytosolic sensor of viral DNA, it has mainly been detected in the nucleus and was shown to interact with nuclear herpes viral DNA (16, 40, 41). The pyrin and HIN domain (PYHIN) containing proteins, such as IFI16, are also known to act as transcriptional regulators. Recent data showed that IFI16 inhibits HCMV transcription (14) and restricts HSV-1 replication by repressing viral gene expression independently of innate immune sensing (15, 42), thereby constituting an innate immune sensor for cytosolic and nuclear ds- as well as ssDNA. AAV2 DNA being present as ss-, ds- and circular dsDNA might therefore provoke an IFI16 triggered reaction. Indeed, the post-transcriptional silencing of *IFI16* increased AAV2 transduction efficiency, regardless of the structure of the vector genome, indicating an IFI16-mediated inhibition of AAV2 vector transduction. This IFI16-mediated inhibition, however, was shown to be Jak1 and STING independent, suggesting an immune-modulatory independent mode of action. Our data, however, indicate a putative interplay of IFI16 and AAV2 gene expression, as the post-transcriptional silencing of *IFI16* increased AAV2 *rep* expression. Several studies showed that IFI16 exerts its effects by various genome regulation mechanisms, independently of innate immune sensing, including the change of methylation marks (15, 16) or by affecting viral gene expression by reducing the availability of the transcription factor Sp1 (19). Although it has long been known that Sp1 plays a key role in the Rep-mediated induction of wild-type AAV promoter regions (43, 44), its relevance in AAV2 biology has hardly been explored because it is commonly accepted that Sp1 is ubiquitously and constitutively expressed. However, our data indicate that IFI16 reduces the availability of Sp1, thereby suppressing the Sp1-mediated activity.

Overall, we present here the first transcriptome analysis of AAV2-infected human primary fibroblasts. The high quality of the raw data allowed a broad and comparative analysis of the transcription profiles of mock-infected and AAV2-infected cells in absence of a helper virus. Most importantly, our findings provide evidence that not only Toll-like receptor 9 (TLR9) detects AAV genomes and triggers an antiviral state upon AAV infection (45, 46), along with the transgenic genome-derived dsRNA-induced MDA5-mediated innate immune response (47), but also additional sensors such as IFI16 constitute other lines of antiviral defense by suppressing viral gene expression in an Sp1-dependent manner. Hence, further studies on the role of other PYHIN proteins as effectors of antiviral defense mechanism in AAV2 infection or AAV vector-mediated cell transduction seem highly warranted.

## MATERIALS AND METHODS

### Cells

Normal human fibroblast (NHF) cells were kindly privded by X.O. Breakefield (Massachusetts General Hospital, Charlestown, MA, USA) and HeLa cells were maintained in growth medium containing Dulbecco’s modified Eagle medium (DMEM) supplemented with 10% fetal bovine serum (FBS), 100 U/ml penicillin G, 100 µg/ml streptomycin, and 0.25 µg/ml amphotericin B (1% AB) at 37°C in a 95% air-5% CO_2_ atmosphere. IFI16 knock-out human bone osteosarcoma epithelial cells (U2OS IFI16^-/-^), as well as the parental cell line (U2OS wt) were kindly provided by Dr. Bala Chandran (Chicago Medical School, RFUMS, USA) and cultured in growth medium containing DMEM supplemented with GlutaMax, 10% FBS, 1% AB at 37°C in a 95% air-5% CO_2_ atmosphere. 2fTGH Jak1^-/-^ (UA4 cell line, 12021505, Sigma-Aldrich, Merck KGaA, Darmstadt, Germany) were maintained growth medium DMEM supplemented with GlutaMax, 10% FBS, 1% AB at 37°C in a 95% air-5% CO_2_ atmosphere.

### Viruses

Wild-type (wt) AAV2 was produced by H. Büning (Hannover Medical School, Hannover, Germany). UV-irradiated AAV2 (UV-AAV2) was produced by UV inactivation of wtAAV2 with 254-nm UV light at a dose of 960 mJ/cm^2^ carried out in a UVC 500 UV cross-linker (Hoefer, Inc., San Francisco, CA, USA). UV-inactivation was assessed on protein level using an anti-Rep antibody (data not shown).

Recombinant (r)AAVeGFP, rAAVmCherry and self-complementary (sc)AAVeGFP vectors of AAV serotype 2 were produced by transient transfection of 293T cells with pDG (48) and pAAVeGFP (kindly provided by M. Linden, King’s College London School of Medicine, London, UK), pAAVmCherry or pscAAVeGFP (kindly provided by J. Neidhardt, University of Zurich, Switzerland), respectively and purified by an iodixanol density gradient. Titers of genome-containing particles were determined by qPCR (49). Lentiviral vectors expressing GFP (TR30021V) or *IFI16* fused to monomeric GFP (RC202193L2V) were obtained from OriGene (Rockville, USA).

### Virus infection for RNA sequencing

5x10^6^ NHF cells were seeded into 10-cm tissue culture plates. The following day, the cells were either mock-infected or infected with AAV2 (multiplicity of infection [MOI], of 500) in DMEM (0% FBS, 1% AB; pre-cooled to 4°C). Virus was allowed to adsorb at 4°C for 30 min before cultures were placed for 1 h into a humidified incubator at 37°C in a 95% air-5% CO_2_ atmosphere. After washing the cells with phosphate-buffered saline (PBS) and adding fresh medium (DMEM supplemented with 2% FBS and 1% AB), the cells were placed back at 37°C in a 95% air-5% CO_2_ atmosphere.

### RNA extraction

Cells were infected as described above. After 24 h, total RNA was extracted using the Direct-zol^TM^ RNA MiniPrep kit according to the instructions of the manufacturer (Zymo Research Corp, Irvine, CA, USA). DNA was digested by adding 8 µl of 10X DNase buffer, 5 µl DNase, 3 µl RNase-free water, and 64 µl RNA wash buffer and incubation for 15 min at 37°C. The samples were then purified according to the manufacturer’s (Zymo Research Corp) protocol. Quality and quantity of the extracted RNA was assessed using Bioanalyzer 2100 (Agilent Technologies, Inc., Santa Clara, CA, USA). Samples with an RNA Integrity Number (RIN) of at least 8.3 were further used for the RNA-seq analyses.

### Illumina RNA Sequencing

The RNA-seq experiment was performed in four steps. (i) a cDNA library was prepared from the RNA, (ii) cDNA was amplified in clusters, (iii) clusters were sequenced and (iv) primary sequencing data were analyzed.

### Library preparation

The Illumina TruSeq Stranded Total RNA Sample Prep Kit with Ribo-Zero Human/Mouse/Rat protocol (Illumina, Inc. San Diego, CA, USA) was used for the following steps: 1 µg of total RNA was freed of cytoplasmic rRNA using biotinylated, Human/Mouse/Rat specific oligonucleotides combined with Ribo-Zero rRNA removal beads and further fragmented into small pieces by divalent cations under elevated temperatures. First strand cDNA was synthesized using Reverse Transcriptase II, Actinomycin D and random primers. Second strand cDNA synthesis was achieved by removing the RNA template and synthesizing a replacement strand, which incorporates dUTP instead of dTTP to generate double stranded cDNA. The resulting cDNA samples were fragmented, 3‘adenylated and ligated to multiple indexing adaptors. Fragments containing those adaptors on both ends were selectively enriched using PCR. The quality and quantity of the enriched libraries were validated using Bioanalyzer 2100 (Agilent Technologies). Diluted libraries (10 nM) were pooled and further used for cluster generation.

### Cluster Generation and Sequencing

The TruSeq SR Cluster Kit v3-cBot-HS (Illumina, Inc.) was used for cluster generation using diluted (10 nM) and pooled libraries. Sequencing was performed on the Illumina HiSeq 2500 in the high throughput mode. Library preparation and sequencing was performed at the Functional Genomics Center Zurich (FGCZ) core facility

### Sequencing data analysis

Reads were aligned with the STAR aligner (STAR: ultrafast universal RNA-seq aligner with the additional parameters (-- outFilterMatchNmin 30 - outFilterMismatchNmax 5 – outFilterMismatchNoverLmax 0.05 -- outFilterMultimapNmax 50), which means that at least 30 bp matching are required and that at most 5 mismatches and 5% of mismatches are accepted. Read alignments were only reported for reads with less than 50 valid alignments. The Human genome build and annotation from Ensembl (GRCh37) was used as reference. Spliced junctions derived from the Ensemble gene annotations. Additionally, the reference was extended to contain the AAV2 sequence (Genebank accession #NC_001401). Expression counts were computed using the R Bioconductor package GenomicRanges (50). Differential expression was computed using the R DESeq2 package (51).

### Bioinformatic analyses

Gene Ontology (GO) term biological process (BP) analysis was performed by DAVID (22). An enrichment map of the DAVID GOterms BP analysis was constructed using the Cytoscape module Enrichment Map. Heat maps of genes representing selected ontologies were constructed using R KEGG pathway analysis (R Bioconductor package Pathview) (52).

### Antibodies

The following primary antibodies were used: anti-β-actin (Sigma-Aldrich A5316; dilution for Western blotting [WB]; 1:10‘000), anti-cyclin A (BD Biosciences; dilution for WB; 1:250), anti-cyclin B1 (Cell Signaling 4138; dilution for WB; 1:1‘000), anti-CDK1 (Abcam; dilution for WB; 1:1‘000), anti-E2F1 (Cell Signaling 3742; dilution for WB; 1:1‘000), anti-p53 (Abcam; dilution for WB; 1:1‘000), anti-Rb (Cell Signaling 9309; dilution for WB; 1:2‘000), anti-Rb-P-S807/811 (Cell Signaling 8516; dilution for WB; 1:1‘000), anti-IFI16 (Santa Cruz Biotechnology 1G7; dilution for WB; 1:500, dilution for immunofluorescence [IF]; 1:250), anti-STING (Santa Cruz Biotechnology E-20; dilution for WB; 1:500), anti-Rep (RDI, Division of Fitzgerald Industries; dilution for WB; 1:200), anti-AAV2 intact particle (A20, ProGen: dilution for IF; 1:50), anti-fibrillarin (Abcam ab5821; dilution for WB; 1:650), anti-Sp1 (Abcam ab227383; ChIP and co-immunoprecipitation [Co-IP] assays; 1.5 μg, dilution for WB; 1:500) or rabbit IgG1 antibody (Abcam ab171870; ChIP and Co-IP assays; 1.5 μg). The following secondary antibodies were used: rabbit-anti mouse IgG-horseradish peroxidase (HRP; SouthernBiotech; dilution 1:10‘000) and goat-anti rabbit IgG-HRP (SouthernBiotech; dilution: 1:10‘000)

### Western blotting

A total of 1.5x10^6^ NHF cells were seeded into 10-cm tissue culture plates. The following day, the cells were either mock-infected or infected with AAV2 (MOI 500) in DMEM (0% FBS, 1% AB; pre-cooled to 4°C). Virus was allowed to adsorb at 4°C for 30 min before cultures were placed for 1 h into a humidified incubator at 37°C in a 95% air-5% CO_2_ atmosphere. After washing the cells with PBS and adding fresh medium (DMEM supplemented with 2% FBS and 1% AB), the cells were placed back at 37°C in a 95% air-5% CO_2_ atmosphere. 48 h later the cells were trypsinized, washed once with PBS, and centrifuged for 5 min at 2000 x g and 4°C. The pellet was dissolved in 100 μl protein loading buffer (2.5% SDS, 5% β-mercaptoethanol, 10 % glycerol, 0.002% bromophenol blue, 62.5 mM Tris-HCl, pH 6.8) and then the samples were boiled for 10 min. Cell lysates were separated, depending on the molecular weight of the protein of interest, on 10% or 12% SDS-polyacrylamide gels and transferred to Protran nitrocellulose membranes (Whatman, Bottmingen, Switzerland). Membranes were blocked with PBS-T (PBS containing 0.3% Tween 20) supplemented with 5% nonfat dry milk for 1 h at room temperature (RT). Incubation with antibodies was carried out with PBS-T supplemented with 2.5% milk. Primary antibodies were incubated overnight at 4°C while secondary antibodies were incubated for 1 h at RT. Membranes were washed 3 times with PBS-T for 10 min after each antibody incubation step. HRP-conjugated secondary antibodies were detected by incubation with ECL (WesternBright^TM^ ECL-spray, Advansta Inc., Menlo Park, CA, USA) for 2 min. The membranes were exposed to chemiluminescence detection films (Roche Diagnistics, Rotkreuz, Switzerland). Detection of anti-actin served as loading control for the lysate.

### Quantitative reverse transcription PCR (RT-qPCR)

For primer design, the Primer-BLAST tool, the Harvard primerbank and the PrimerCheck of the SpliceCenter was used. To test the primers, a standard RT-PCR (+/- RT reaction) with mock-infected NHF cells was performed. The cycling protocol started with a denaturation step of 3 min at 95°C, followed by 37 cycles of 30 sec at 94°C, 30 sec at 55°C and 1 min at 72°C followed by a final step of 10 min at 72°C. Subsequently, the reactions were analyzed on 1% agarose gels. Bands were expected between 100-200 bp, depending on the primer pair. The concentration and purity of RNA was determined by Qubit fluorometer analysis. To generate cDNA, the extracted RNA was reverse transcribed by using the reverse transcription system (Promega Corporation, Fitchburg, WI, USA). For this, the following components were mixed: 4 µl MgCl_2_, 2 µl 10X RT buffer, 2 µl dNTPs (10 mM), 0.5 µl of the RNase inhibitor RNAsin, 0.65 µl AMV RT, 1 µl random or Oligo(dT) primers, 1 µg RNA and RNase free H_2_O in a total volume of 20 µl. The mixture was incubated for 10 min at room temperature and 15 min at 42°C. For enzyme inactivation, the sample was incubated for 5 min at 95°C and then incubated on ice for 5 min. 4 µl of the cDNA (approximately 10 ng) were used for qPCR and the rest was stored at -20°C. For each reaction the following mixture was prepared: 1 μl forward primer (10 μM), 1 μl reverse primer (10 μM), 10 µl of SYBR Green PCR master mix and 4 µl ddH_2_O and transferred into a well of a Hard-Shell 96-well PCR plate (MicroAmp fast 96-well reaction plate). 4 µl of the appropriate cDNA was added and the 96-well plate was centrifuged for 1 min at 1000 x g and subsequently run at the standard 20 µl qPCR SYBR green program on QuantStudio 3 real-time system (Applied Biosystem, ThermoFisher Scientific, Waltham, MA, USA). The experiment was performed as technical triplicates for each primer pair for infected and non-infected samples. GAPDH and SDHA were used as housekeeping genes for the further normalization of the RT-qPCR raw data. Used primer sequences are listed in Table 2.

**TABLE 2.**
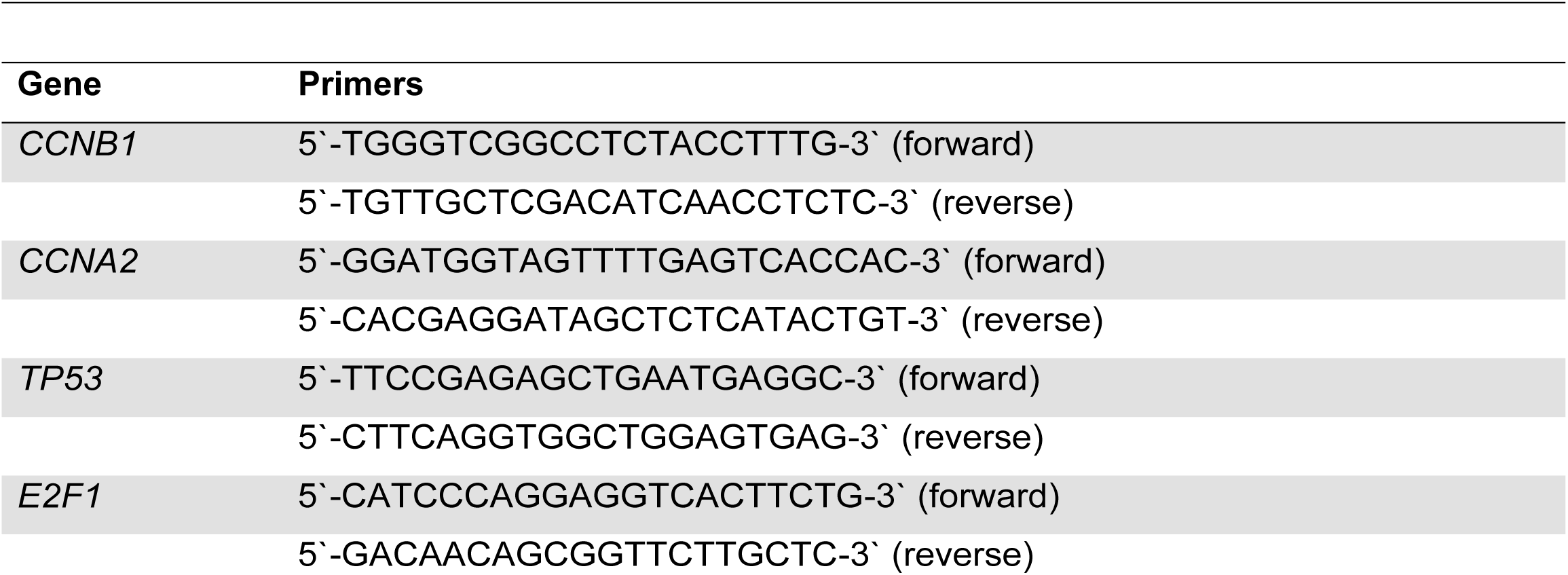
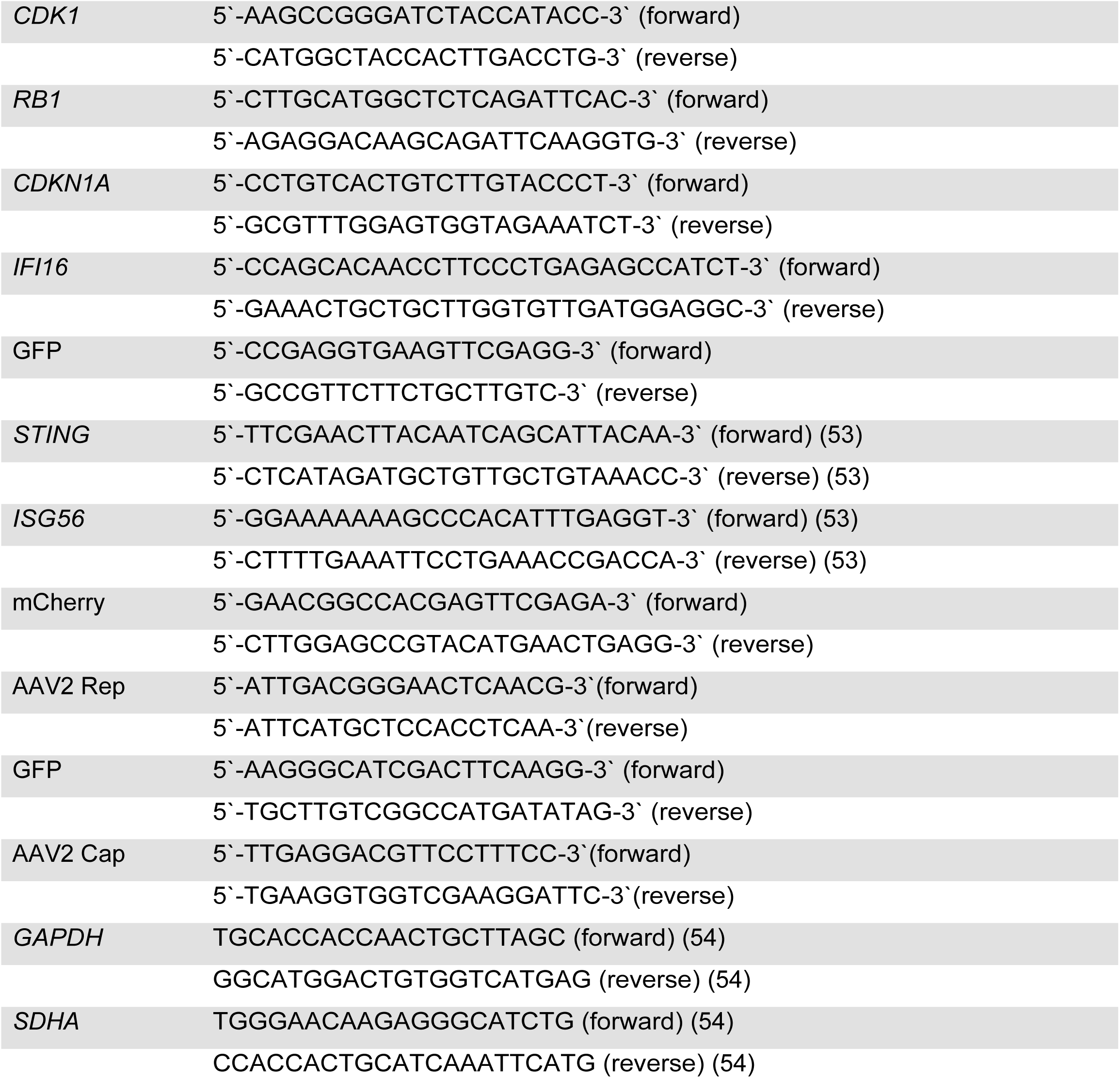
RT-qPCR primers used in this study.

### Quantitative PCR (qPCR)

For qPCR, total DNA was isolated by using the DNeasy blood and tissue kit (Qiagen, Hilden, Germany) according to the manufacturer’s protocol. 4 µl of the isolated DNA (approximately 10 ng) were used for qPCR and the rest was stored at -20°C. For each reaction the following mixture was prepared: 1 μl forward primer (10 μM), 1 μl reverse primer (10 μM), 10 µl of SYBR Green PCR master mix and 4 µl ddH_2_O and transferred into a well of a Hard-Shell 96-well PCR plate (MicroAmp fast 96-well reaction plate). 4 µl of the appropriate DNA was added and the 96-well plate was centrifuged for 1 min at 1000 x g and subsequently run at the standard 20 µl qPCR SYBR green program on QuantStudio 3 real-time system (Applied Biosystem, ThermoFisher Scientific, Waltham, MA, USA). The experiment was performed as technical triplicates for each primer pair for infected and non-infected samples. The transcriptional start site (TSS) of GAPDH was used as endogenous control. Used primer sequences are listed in Table 3.

**Table 3.**
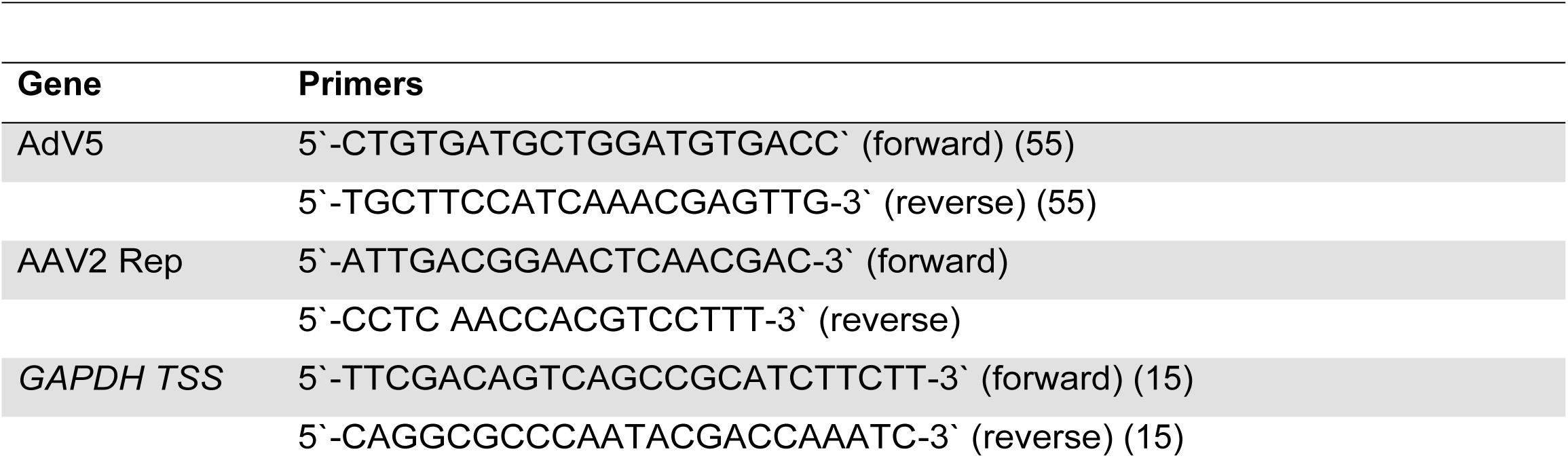
qPCR primers used in this study.

### Co-immunoprecipitation (Co-IP)

1.3x10^6^ U2OS IFI16^-/-^ or U2OS wt were washed twice with cold PBS and harvested using a cell scraper, and transferred into a 15 ml conical tube while being kept on ice. Next, the cells were pelleted at 900 g for 10 min at 4°C. The remaining pellet was resuspended in 50 μl PBS and 100 μl of 2X SDS lysis buffer (100 mM Tris-HCl, pH 8.1, 2% SDS (w/v), 20 mM EDTA) with protease inhibitor (cOmplete mini, Roche Cat. 11836153001) was added. After 15 min of incubation on ice, the samples were centrifuged again, and the lysate was diluted 1:10 in dilution buffer (16.7 mM Tris-HCL, pH 8.1, 167 mM NaCl, 0.01% SDS (w/v), 1.2 mM EDTA, 1.1% Triton X-100 (v/v)). Next, 80 μl agarose protein G with salmon sperm DNA slurry (Millipore Cat. 16-201) was added for pre-clearance, and the samples were incubated with gentle agitation at 4°C for 30 min. The slurry was then centrifuged for 3 min, 4°C at 300 g (to not break the agarose beads), and the supernatant was recovered and split into two equal fractions. The fractions were then incubated either with 1.5 μg of anti-Sp1 antibody or rabbit IgG1 antibody overnight at 4°C with gentle agitation. The next day, 120 μl agarose protein G with salmon sperm DNA slurry was added and incubated for 30 min at 4°C. The centrifugation for the following washing steps were performed at 4 °C at 300 g for 3 min each. First, the samples were washed twice with low salt wash buffer (20 mM Tris-HCl, pH 8.1, 150 mM NaCl, 0.1% SDS (w/v), 2 mM EDTA, 1% Triton X-100 (v/v)) and then with high salt wash buffer (20 mM Tris-HCl, pH 8.1, 500 mM NaCl, 0.1% SDS (w/v), 2 mM EDTA, 1% Triton X-100 (v/v)). Next, the samples were washed with LiCl salt wash buffer (20 mM Tris-HCl, pH 8.1, 250 mM LiCl, 1% deoxycholate (w/v), 1 mM EDTA, 1% Nonidet P-40 (v/v)), and finally twice with TE (pH 8.0). After centrifugation, the supernatant was discarded and the samples were dissolved in 6X protein loading buffer, boiled for 10 min, and subjected to Western blot analysis.

### Chromatin immunoprecipitation (ChIP)

5x10^5^ U2OS IFI16^-/-^ or U2OS wt were either untransduced, or transduced with lentiviral vectors expressing either GFP (MOI 5) or *IFI16* fused to monomeric GFP (MOI 5) in presence of polybrene (8 μg/ml, Pierce, Rockford, IL). 72 hours later, 70% confluent T-150 cell culture flasks were either mock-infected or infected with AAV2 (MOI 20‘000), rAAVeGFP (MOI 20‘000), or scAAVeGFP (MOI 20‘000). 24 hpi, the cells were washed once with ice-cold PBS, cross-linked with 1% formaldehyde in PBS, and incubated in a humidified, 95% air- and 5% CO_2_-incubator at 37°C for 10 min. In order to stop the cross-linking, 125 mM glycine was added and the cells were incubated for 5 min at RT. Next, the cells were washed twice with PBS, harvested using a cell scraper, and transferred into a 15 ml conical tube while being kept on ice. Next, the cells were pelleted at 1000 g for 10 min at 4°C. The remaining pellet was resuspended in 50 μl PBS and 100 μl of 1X SDS lysis buffer (100 mM Tris-HCl, pH 8.1, 1% SDS (w/v), 20 mM EDTA) containing protease inhibitor (cOmplete mini, Roche Cat. 11836153001). After 15 min of incubation on ice, the samples were centrifuged again, sonicated (100% amplitude, 15 sec on, 15 sec off, 20 min total sonication time), and the lysate was diluted 1:10 in dilution buffer (16.7 mM Tris-HCL, pH 8.1, 167 mM NaCl, 0.01% SDS (w/v), 1.2 mM EDTA, 1.1% Triton X-100 (v/v)). Next, 80 μl agarose protein G with salmon sperm DNA slurry (Millipore Cat. 16-201) was added for pre-clearance, and the samples were incubated with gentle agitation at 4°C for 30 min. The slurry was then centrifuged for 3 min, 4°C at 300 g (to not break the agarose beads), and the supernatant was recovered and splitted into two equal fractions. The fractions were then incubated either with 1.5 μg of anti-Sp1 antibody or rabbit IgG1 antibody at 4°C overnight with gentle agitation. The next day, 120 μl agarose protein G with salmon sperm DNA slurry was added and incubated for 30 min at 4°C. The centrifugation for the following washing steps were performed at 4 °C at 300 g for 3 min each. First, the samples were washed twice with low salt wash buffer (20 mM Tris-HCl, pH 8.1, 150 mM NaCl, 0.1% SDS (w/v), 2 mM EDTA, 1% Triton X-100 (v/v)) and then with high salt wash buffer (20 mM Tris-HCl, pH 8.1, 500 mM NaCl, 0.1% SDS (w/v), 2 mM EDTA, 1% Triton X-100 (v/v)). Next the samples were washed with LiCl salt wash buffer (20 mM Tris-HCl, pH 8.1, 250 mM LiCl, 1% deoxycholate (w/v), 1 mM EDTA, 1% Nonidet P-40 (v/v)), and finally twice with TE (pH 8.0). After centrifugation, the supernatant was discarded, and the samples were eluted twice in 250 µl of freshly prepared elution buffer (1% SDS (w/v), 100 mM NaHCO_3_) for 15 min at 65°C, and eluates were combined. Next, 20 μl 5 M NaCl were added to the samples and they were incubated at 65°C overnight in order to reverse the cross-link. To recover the DNA, 10 μl 0.5 M EDTA, 20 μl 1 M Tris-HCl (pH 6.5), 2 μl proteinase K (10 mg/ml), and 1 μl RNase A (10 mg/ml) were added, and the samples were incubated at 45°C for 1 h. For phenol/chloroform extraction of the DNA, 1 volume of phenol:chloroform:isoamylalcohol (25:24:1, v/v, 15593031, Invitrogen, USA) was added. The samples were centrifuged for 5 min at 4°C and 15’500 g. The supernatant was transferred into a fresh tube, and 1 volume chloroform was added. The sample was centrifuged for 1 min at 4°C and 15’500 g, and the supernatant was transferred into a fresh tube. 2.5 volumes of EtOH (pure), and 0.1 volume of 3M NaAc (pH 5.5) were added to the sample. To precipitate the DNA the suspension was incubated for at least 20 min at -80°C. Next, the samples were centrifuged for 10 min at 4°C and 18’000 g, and the supernatant was discarded. The DNA pellet was washed with 70% EtOH and centrifuged for 10 min at 4°C and 18’000 g. The supernatant was removed, and the DNA pellet was left to dry for at least 20 min at RT. After drying, the pellet was resuspended in Tris-HCl (pH 8.5) and incubated for 10 min at 37°C. For qPCR, 4 µl of the isolated DNA (approximately 10 ng) were used, and the rest was stored at -20°C. For each reaction the following mixture was prepared: 1 μl forward primer (10 μM), 1 μl reverse primer (10 μM), 10 µl of SYBR Green PCR master mix and 4 µl ddH_2_O and transferred into a well of a Hard-Shell 96-well PCR plate (MicroAmp fast 96-well reaction plate). 4 µl of the appropriate DNA was added, and the 96-well plate was centrifuged for 1 min at 1000 x g and subsequently run at the standard 20 µl qPCR SYBR green program on QuantStudio 3 real-time system (Applied Biosystem, ThermoFisher Scientific, Waltham, MA, USA). The experiment was performed as technical triplicates for each primer pair for each sample. Used primer sequences are listed in Table 4.

**Table 4.**
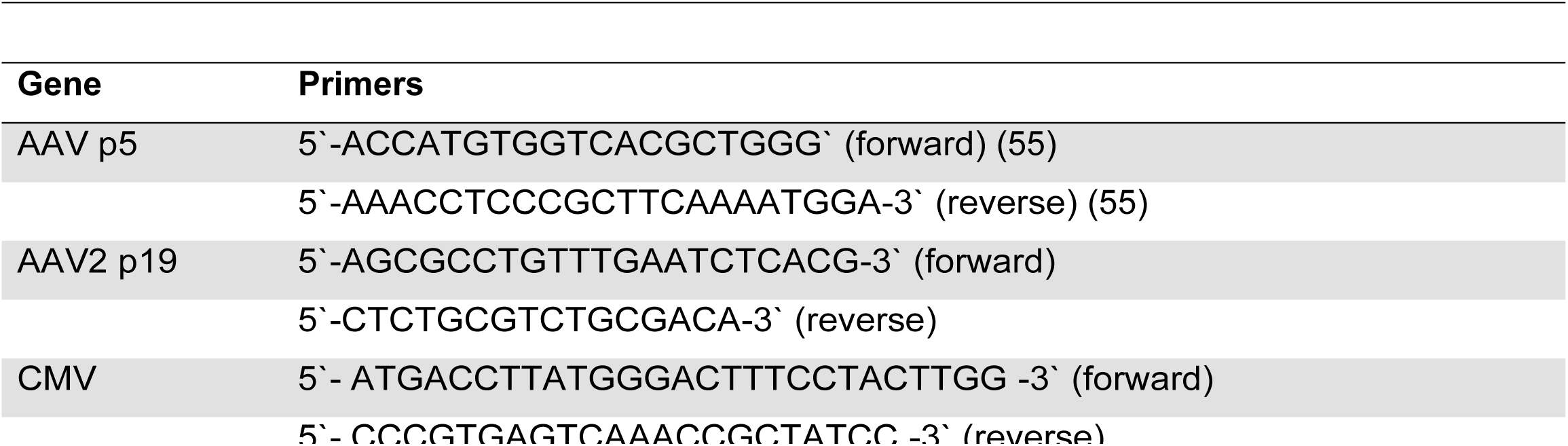
ChIP primers used in this study.

### Cell cycle analysis

NHF cells were seeded onto 6 well tissue culture plates at a confluency of 30% per plate and 24 h later infected with AAV2 or rAAVeGFP (MOI 5‘000) in DMEM (0% FBS, 1% AB; pre-cooled to 4°C). Virus was allowed to adsorb at 4°C for 30 min before cultures were placed for 1 h into a humidified incubator at 37°C in a 95% air-5% CO_2_ atmosphere. After washing the cells with PBS and adding fresh medium (DMEM supplemented with 2% FBS and 1% AB), the cells were placed back at 37°C in a 95% air-5% CO_2_ atmosphere. At the indicated time points the cells were harvested by exposing them to 0.05% Trypsin-EDTA solution for 10 min, centrifuged and washed with PBS, fixed in 2.5 mL ice-cold 100% ethanol and stored overnight at −20°C. At the time of analysis, the cells were centrifuged, washed once again with PBS and stained with a freshly made solution containing 0.1 mg/mL propidium iodide (PI), 0.05% Triton X-100 and 0.1 mg/mL ribonuclease A (RNase A) in PBS. All samples were incubated for 40 min at 37°C in the dark. Cell cycle distribution was determined by flow cytometry (Gallios flow cytometer; Beckman Coulture, Brea, CA, USA), and data was analyzed by using Kaluza Flow Analysis software (Beckman Coulter, Brea, CA, USA).

### RNA interference

6×10^4^ NHF cells per well were plated in 24-well tissue culture plates and transfected using lipofectamine RNAiMax transfection reagent (ThermoFisher scientific, Waltham, MA, USA) according to the manufacturer’s recommendations. The sequences of the siRNAs specific for *IFI16* (pool, 5‘-UTR and cds), C911 siRNA controls and siRNA targeting the cds of STING are listed in Table 5. 40 hpt the cells were transduced with recombinant AAV2 vectors as indicated in the results and figure legends. Knock-down efficiency was assessed either by Western blotting or RT-qPCR.

**Table 5.**
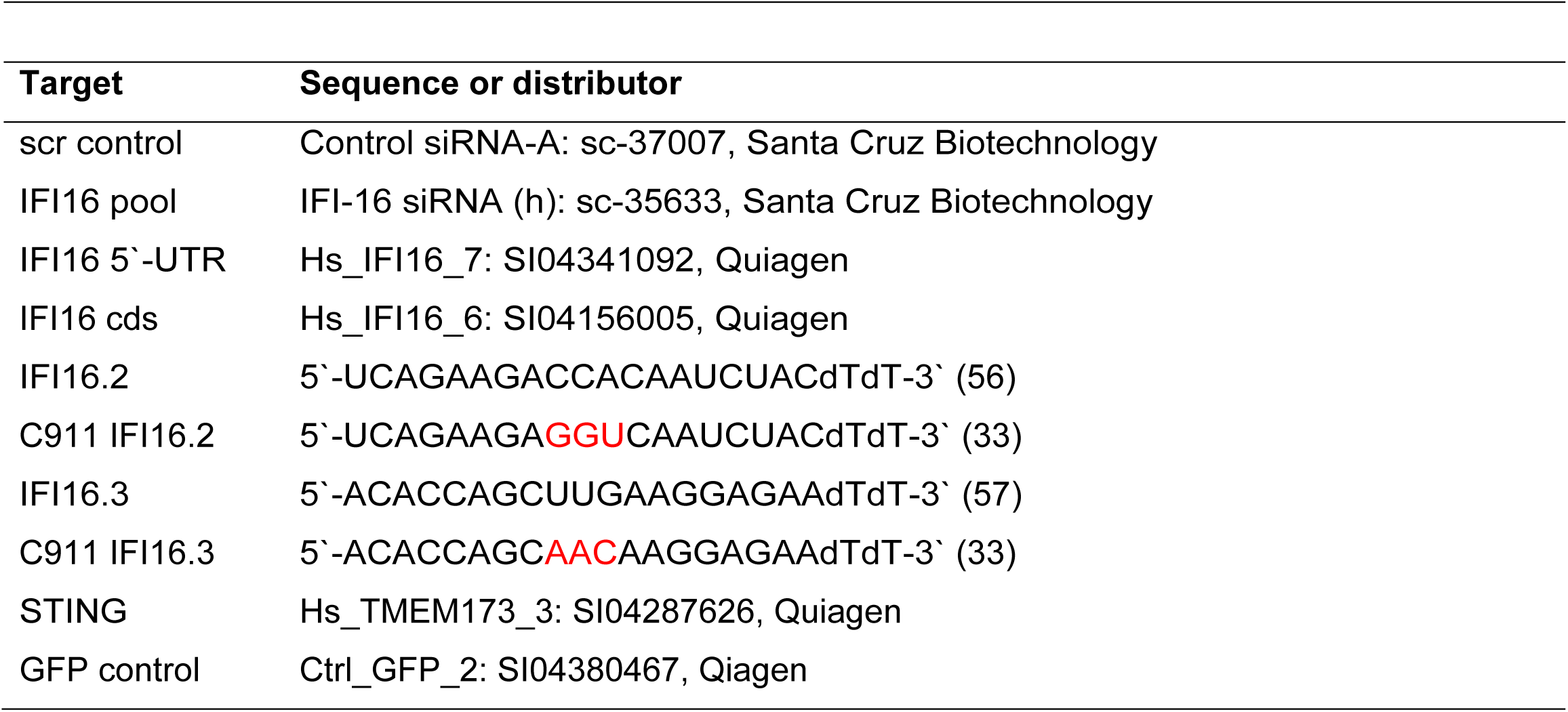
siRNAs used in this study.

### Exogenous complementation

6×10^4^ U2OS IFI16^-/-^ were either untransduced, transduced with lentiviral vectors expressing GFP (MOI 5), or lentiviral vectors expressing *IFI16* fused to monomeric GFP (MOI 5) in presence of polybrene (8 μg/ml, Pierce, Rockford, IL). 72 hours later, the cells were infected with rAAV2mCherry (MOI 500) and, 24 hpi, mCherry and *IFI16* expression was assessed by RT-qPCR using specific primers for mCherry or *IFI16* (Table 1), respectively.

### Microscopy

NHF cells were seeded onto coverslips (12-mm diameter; Glaswarenfabrik Karl Hecht GmbH & Co. KG, Sondheim, Germany) in 24-well tissue culture plates (4x10^4^ cells per well). The next day, the cells were infected as indicated in the results and the figure legends. For immunofluorescence analysis and CSLM, the cells were washed once with cold PBS 24 h after infection and then fixed with 2% paraformaldehyde (PFA) in PBS for 10 min at RT. The fixation process was stopped by incubation with 0.1 M glycine for 10 min at RT and two washes with cold PBS. Afterwards, the cells were permeabilized with 0.1% Triton-X 100 (in PBS) for 10 min, followed by 3 washing steps with PBS. The cells were blocked for 30 min with 3% bovine serum albumin (BSA) in PBS-T (0.05% Tween) at 4°C. For staining, the cells were incubated with antibodies diluted in PBS-T-BSA (3%) in a humidified chamber at RT in the dark. The coverslips were placed onto droplets (20 μl) of a primary or secondary antibody solution. After incubation for 1 h, the cells were washed three times with PBS and once with H_2_O. All coverslips were embedded in ProLong Anti-Fade mountant (Molecular Probes) and cells were observed by using a confocal laser scanning microscope (Leica SP8; Leica Microsystems, Wetzlar, Germany). To prevent cross talk between the channels for the different fluorochromes, all channels were recorded separately, and fluorochromes with longer wavelengths were recorded first.

### Fluorescence *in situ* hybridization

FISH was performed essentially as described previously by Lux et al. (58). Briefly, a 3.9-kb DNA fragment containing the AAV2 genome without the inverted terminal repeats was amplified by PCR from plasmid pDG using forward (5‘-CGGGGTTTTACGAGATTGTG-3‘) and reverse (5‘-GGCTCTGAATACACGCCATT-3‘) primers and the following conditions: 30 s at 95°C; 35 cycles of 10 s at 98°C, 15 s at 58°C, and 75 s at 72°C; and 10 min at 72°C. The PCR sample was then digested with DpnI to cut the residual template DNA and purified with the Pure Link PCR purification kit (Qiagen, Hilden, Germany). The DNA fragment was labeled with 5-(3-aminoallyl)dUTP by nick translation, and the incorporated dUTPs were labeled with amino-reactive Alexa Fluor 647 dye by using the Ares DNA labeling kit (Molecular Probes, Eugene, OR, USA) according to the manufacturer’s protocols. NHF cells were plated onto glass coverslips in 24-well plates at a density of 4x10^4^ cells per well and 24 h later, the cells were mock infected or infected with AAV2 (MOI of 20‘000). 24 hours after infection, the cells were washed with PBS, fixed for 30 min at RT with 2% PFA (in PBS), and then washed again with PBS. The cells were then quenched for 10 min with 50 mM NH_4_Cl (in PBS), washed with PBS, permeabilized for 10 min with 0.2% Triton X-100 (in PBS), blocked for 10 min with 0.2% gelatin (in PBS), and washed again with PBS. Hybridization solution (20 μl per coverslip) containing 1 ng/μl of the labeled DNA probe, 50% formamide, 7.3% dextran sulfate, 15 ng/μl salmon sperm DNA, and 0.74x SSC (1x SSC is 0.15 M NaCl plus 0.015 M sodium citrate) was denatured for 3 min at 95°C and shock-cooled on ice. The coverslips with the fixed, and permeabilized cells facing down were placed onto a drop (20 μl) of the denatured hybridization solution and incubated overnight at 37°C in a humidified chamber (note that the cells were not denatured, as the AAV2 genome is present as ssDNA). The next day, the coverslips were washed three times with 2x SSC at 37°C, three times with 0.1x SSC at 60°C, and twice with PBS at RT. The cells were then embedded in ProLong Anti-Fade mountant (Molecular Probes, Eugene, OR, USA) and imaged by confocal laser scanning microscopy (Leica SP8; Leica Microsystems, Wetzlar, Germany).

### Data availability

Sequencing data (full length and p < 0.01, number of reads > 40) are available under Zenodo (doi.org/10.5281/zenodo.7147541).

## ACKNOWLEDGMENTS

We would like to thank Hubert Rehrauer (Functional Genomics Center Zurich, University of Zurich) for the support with RNA sequencing and for assistance with data analysis.

This work was supported by a grant (310030_184766) from the Swiss National Science Foundation to C.F.

## Legends Supplementary Figures

**FIG S1** Selected heat maps of affected biological processes. Reads of the 50 most differentially expressed genes (depicted on the right of each heat map) between AAV2-infected cells (A1-A3) and mock-infected cells (M1-M3) from (A) the cell cycle and (B) chromatin organization clusters of the enrichment map. The dendrogram (left side of each heat map) illustrates the unsupervised clustering of the genes.

**FIG S2** Clustered heat map of 872 genes with a log_2_ ratio > |0.58| and a significance threshold of p < 0.01 were used for the unsupervised clustering of the genes. AAV2-infected samples (A1-A3) are represented on the left and mock-infected samples (M1-M3) on the right.

**FIG S3** Cell cycle phase distributions upon AAV2 or rAAV2 infection over time. NHF cells were infected with AAV2 or rAAVeGFP (MOI 5‘000). At the indicated time points, the cell cycle profile was assessed by PI staining and flow cytometry (10‘000 cells per sample). Graph shows mean and SD of the percentage of cells in each cell cycle phase at the individual time points. p-values were calculated using an unpaired Student‘s t-test (* - p ≤ 0.05, ** - p ≤ 0.01, *** - p ≤ 0.001, **** - p ≤ 0.0001).

**FIG S4** C911 siRNA controls. NHF cells were reverse transfected with no siRNA, scr control siRNA or siRNAs targeting the coding sequence of *IFI16* (IFI16 pool, IFI16.2 and IFI16.3). To address the question of off-target effects of the individual siRNAs (IFI16.2, IFI16.3), C911 siRNA controls were included. 36 hpt the cells were either mock-infected or infected with rAAVeGFP (MOI 4‘000). (A) 24 hpi, cells were counted using a fluorescence microscope. (B) The graph shows mean and SD of the relative cell count of GFP positive NHF cells from triplicate experiments. p-values were calculated using an unpaired Student‘s t-test (* - p ≤ 0.05, ** - p ≤ 0.01, *** - p ≤ 0.001, **** - p ≤ 0.0001). (C) Knock-down of *IFI16* was confirmed on protein level.

**FIG S5** STING signaling in different cell lines. NHF, U2OS and HeLa cells were treated with 2’3’-cGAMP (3 μM) for 9 h, and total RNA was extracted, converted to cDNA and subjected to RT-qPCR using specific primers for *STING* and *ISG56*. p-values were calculated using an unpaired Student‘s t-test (* - p ≤ 0.05, ** - p ≤ 0.01, *** - p ≤ 0.001, **** - p ≤ 0.0001).

**FIG S6** Nucleolar localization of IFI16. NHF cells were infected with AAV2 (MOI 20‘000). After 24 h, the cells were fixed and processed for multicolor IF and CLSM. IFI16 was detected by using a monoclonal antibody against IFI16 (green/blue). Nucleoli were visualized using an antibody against fibrillarin (red). Nuclei were counterstained with DAPI.

## REFERENCES

1. Kuzmin DA, Shutova MV, Johnston NR, Smith OP, Fedorin VV, Kukushkin YS, Loo JCM van der, Johnstone EC. 2021. The clinical landscape for AAV gene therapies. Nat Rev Drug Discov 20:173–174.

2. Samulski RJ, Zhu X, Xiao X, Brook JD, Housman DE, Epstein N, Hunter LA. 1991. Targeted integration of adeno-associated virus (AAV) into human chromosome 19. The EMBO journal 10:3941–3950.

3. Sun X, Lu Y, Bish LT, Calcedo R, Wilson JM, Gao G. 2010. Molecular Analysis of Vector Genome Structures After Liver Transduction by Conventional and Self-Complementary Adeno-Associated Viral Serotype Vectors in Murine and Nonhuman Primate Models. Hum Gene Ther 21:750–761.

4. Buller RML, Janik JE, Sebring ED, Rose JA. 1981. Herpes Simplex Virus Types 1 and 2 Completely Help Adenovirus-Associated Virus Replication. J Virol 40:241– 247.

5. Srivastava A, Lusby EW, Berns KI. 1983. Nucleotide sequence and organization of the adeno-associated virus 2 genome. J Virol 45:555–564.

6. Laughlin CA, Westphal H, Carter BJ. 1979. Spliced adenovirus-associated virus RNA. Proc National Acad Sci 76:5567–5571.

7. Pereira DJ, McCarty DM, Muzyczka N. 1997. The adeno-associated virus (AAV) Rep protein acts as both a repressor and an activator to regulate AAV transcription during a productive infection. J Virol 71:1079–1088.

8. Sonntag F, Köther K, Schmidt K, Weghofer M, Raupp C, Nieto K, Kuck A, Gerlach B, Böttcher B, Müller OJ, Lux K, Hörer M, Kleinschmidt JA. 2011. The Assembly-Activating Protein Promotes Capsid Assembly of Different Adeno-Associated Virus Serotypes. J Virol 85:12686–12697.

9. Ogden PJ, Kelsic ED, Sinai S, Church GM. 2019. Comprehensive AAV capsid fitness landscape reveals a viral gene and enables machine-guided design. Science 366:1139–1143.

10. Chu Y, Corey DR. 2012. RNA sequencing: platform selection, experimental design, and data interpretation. Nucleic acid therapeutics 22:271–274.

11. Maher CA, Kumar-Sinha C, Cao X, Kalyana-Sundaram S, Han B, Jing X, Sam L, Barrette T, Palanisamy N, Chinnaiyan AM. 2009. Transcriptome sequencing to detect gene fusions in cancer. Nature 458:97–101.

12. Ingolia NT, Brar GA, Rouskin S, McGeachy AM, Weissman JS. 2012. The ribosome profiling strategy for monitoring translation in vivo by deep sequencing of ribosome-protected mRNA fragments. Nature protocols 7:1534–1550.

13. Unterholzner L, Keating SE, Baran M, Horan KA, Jensen SB, Sharma S, Sirois CM, Jin T, Latz E, Xiao TS, Fitzgerald KA, Paludan SR, Bowie AG. 2010. IFI16 is an innate immune sensor for intracellular DNA. Nature Immunology 11:997–1004.

14. Gariano GR, Dell’Oste V, Bronzini M, Gatti D, Luganini A, Andrea MD, Gribaudo G, Gariglio M, Landolfo S. 2012. The intracellular DNA sensor IFI16 gene acts as restriction factor for human cytomegalovirus replication. PLoS pathogens 8:e1002498.

15. Johnson KE, Bottero V, Flaherty S, Dutta S, Singh VV, Chandran B. 2014. IFI16 restricts HSV-1 replication by accumulating on the hsv-1 genome, repressing HSV-1 gene expression, and directly or indirectly modulating histone modifications. PLoS pathogens 10:e1004503.

16. Orzalli MH, Conwell SE, Berrios C, DeCaprio JA, Knipe DM. 2013. Nuclear interferon-inducible protein 16 promotes silencing of herpesviral and transfected DNA. Proceedings of the National Academy of Sciences of the United States of America 110:E4492–501.

17. Cigno IL, Andrea MD, Borgogna C, Albertini S, Landini MM, Peretti A, Johnson KE, Chandran B, Landolfo S, Gariglio M. 2015. The Nuclear DNA Sensor IFI16 Acts as a Restriction Factor for Human Papillomavirus Replication through Epigenetic Modifications of the Viral Promoters. J Virol 89:7506–7520.

18. McLaren PJ, Gawanbacht A, Pyndiah N, Krapp C, Hotter D, Kluge SF, Götz N, Heilmann J, Mack K, Sauter D, Thompson D, Perreaud J, Rausell A, Munoz M, Ciuffi A, Kirchhoff F, Telenti A. 2015. Identification of potential HIV restriction factors by combining evolutionary genomic signatures with functional analyses. Retrovirology 12:41.

19. Hotter D, Bosso M, Jønsson KL, Krapp C, Stürzel CM, Das A, Littwitz-Salomon E, Berkhout B, Russ A, Wittmann S, Gramberg T, Zheng Y, Martins LJ, Planelles V, Jakobsen MR, Hahn BH, Dittmer U, Sauter D, Kirchhoff F. 2019. IFI16 Targets the Transcription Factor Sp1 to Suppress HIV-1 Transcription and Latency Reactivation. Cell Host Microbe 25:858–872.e13.

20. Swonger JM, Liu JS, Ivey MJ, Tallquist MD. 2016. Genetic tools for identifying and manipulating fibroblasts in the mouse. Differentiation 92:66–83.

21. Davidson S, Coles M, Thomas T, Kollias G, Ludewig B, Turley S, Brenner M, Buckley CD. 2021. Fibroblasts as immune regulators in infection, inflammation and cancer. Nat Rev Immunol 21:704–717.

22. Huang DW, Sherman BT, Lempicki RA. 2009. Systematic and integrative analysis of large gene lists using DAVID bioinformatics resources. Nature protocols 4:44–57.

23. Khatri P, Drăghici S. 2005. Ontological analysis of gene expression data: current tools, limitations, and open problems. Bioinformatics (Oxford, England) 21:3587– 3595.

24. Ogata H, Goto S, Sato K, Fujibuchi W, Bono H, Kanehisa M. 1999. KEGG: Kyoto Encyclopedia of Genes and Genomes. Nucleic acids research 27:29–34.

25. Fussenegger M, Bailey JE. 1998. Molecular regulation of cell-cycle progression and apoptosis in mammalian cells: implications for biotechnology. Biotechnology progress 14:807–833.

26. Vermeulen K, Bockstaele DRV, Berneman ZN. 2003. The cell cycle: a review of regulation, deregulation and therapeutic targets in cancer. Cell Proliferat 36:131– 149.

27. Nagano K, Itagaki C, Izumi T, Nunomura K, Soda Y, Tani K, Takahashi N, Takenawa T, Isobe T. 2006. Rb plays a role in survival of Abl-dependent human tumor cells as a downstream effector of Abl tyrosine kinase. Oncogene 25:493–502.

28. Agarwal ML, Agarwal A, Taylor WR, Stark GR. 1995. p53 controls both the G2/M and the G1 cell cycle checkpoints and mediates reversible growth arrest in human fibroblasts. Proc National Acad Sci 92:8493–8497.

29. Hermeking H, Lengauer C, Polyak K, He T-C, Zhang L, Thiagalingam S, Kinzler KW, Vogelstein B. 1997. 14-3-3σ Is a p53-Regulated Inhibitor of G2/M Progression. Mol Cell 1:3–11.

30. Franzoso FD, Seyffert M, Vogel R, Yakimovich A, Pereira B de A, Meier AF, Sutter SO, Tobler K, Vogt B, Greber UF, Büning H, Ackermann M, Fraefel C. 2017. Cell Cycle-Dependent Expression of Adeno-Associated Virus 2 (AAV2) Rep in Coinfections with Herpes Simplex Virus 1 (HSV-1) Gives Rise to a Mosaic of Cells Replicating either AAV2 or HSV-1. Journal of Virology 91.

31. Roukos V, Pegoraro G, Voss TC, Misteli T. 2015. Cell cycle staging of individual cells by fluorescence microscopy. Nat Protoc 10:334–348.

32. Sutter SO, Lkharrazi A, Schraner EM, Michaelsen K, Meier AF, Marx J, Vogt B, Büning H, Fraefel C. 2022. Adeno-associated virus type 2 (AAV2) uncoating is a stepwise process and is linked to structural reorganization of the nucleolus. Plos Pathog 18:e1010187.

33. Buehler E, Chen Y-C, Martin S. 2012. C911: A bench-level control for sequence specific siRNA off-target effects. PLoS ONE 7:e51942.

34. Li Z, Cai S, Sun Y, Li L, Ding S, Wang X. 2020. When STING Meets Viruses: Sensing, Trafficking and Response. Front Immunol 11:2064.

35. Christensen MH, Paludan SR. 2017. Viral evasion of DNA-stimulated innate immune responses. Cell Mol Immunol 14:4–13.

36. Hermanns J, Schulze A, Jansen-Db1urr P, Kleinschmidt JA, Schmidt R, Hausen H zur. 1997. Infection of primary cells by adeno-associated virus type 2 results in a modulation of cell cycle-regulating proteins. Journal of Virology 71:6020–6027.

37. Jakobsen MR, Bak RO, Andersen A, Berg RK, Jensen SB, Tengchuan J, Jin T, Laustsen A, Hansen K, Ostergaard L, Fitzgerald KA, Xiao TS, Mikkelsen JG, Mogensen TH, Paludan SR. 2013. IFI16 senses DNA forms of the lentiviral replication cycle and controls HIV-1 replication. Proc National Acad Sci 110:E4571– E4580.

38. Jønsson KL, Laustsen A, Krapp C, Skipper KA, Thavachelvam K, Hotter D, Egedal JH, Kjolby M, Mohammadi P, Prabakaran T, Sørensen LK, Sun C, Jensen SB, Holm CK, Lebbink RJ, Johannsen M, Nyegaard M, Mikkelsen JG, Kirchhoff F, Paludan SR, Jakobsen MR. 2017. IFI16 is required for DNA sensing in human macrophages by promoting production and function of cGAMP. Nat Commun 8:14391.

39. Monroe KM, Yang Z, Johnson JR, Geng X, Doitsh G, Krogan NJ, Greene WC. 2014. IFI16 DNA Sensor Is Required for Death of Lymphoid CD4 T Cells Abortively Infected with HIV. Science 343:428–432.

40. Kerur N, Veettil MV, Sharma-Walia N, Bottero V, Sadagopan S, Otageri P, Chandran B. 2011. IFI16 Acts as a Nuclear Pathogen Sensor to Induce the Inflammasome in Response to Kaposi Sarcoma-Associated Herpesvirus Infection. Cell Host Microbe 9:363–375.

41. Li T, Diner BA, Chen J, Cristea IM. 2012. Acetylation modulates cellular distribution and DNA sensing ability of interferon-inducible protein IFI16. Proceedings of the National Academy of Sciences of the United States of America 109:10558–10563.

42. Diner BA, Lum KK, Javitt A, Cristea IM. 2015. Interactions of the Antiviral Factor Interferon Gamma-Inducible Protein 16 (IFI16) Mediate Immune Signaling and Herpes Simplex Virus-1 Immunosuppression. Molecular & cellular proteomics : MCP 14:2341–2356.

43. Pereira DJ, Muzyczka N. 1997. The adeno-associated virus type 2 p40 promoter requires a proximal Sp1 interaction and a p19 CArG-like element to facilitate Rep transactivation. J Virol 71:4300–4309.

44. Pereira DJ, Muzyczka N. 1997. The cellular transcription factor SP1 and an unknown cellular protein are required to mediate Rep protein activation of the adeno-associated virus p19 promoter. J Virol 71:1747–1756.

45. Laredj LN, Beard P. 2011. Adeno-Associated Virus Activates an Innate Immune Response in Normal Human Cells but Not in Osteosarcoma Cells. J Virol 85:13133– 13143.

46. Ashley SN, Somanathan S, Giles AR, Wilson JM. 2019. TLR9 signaling mediates adaptive immunity following systemic AAV gene therapy. Cell Immunol 346:103997.

47. Shao W, Earley LF, Chai Z, Chen X, Sun J, He T, Deng M, Hirsch ML, Ting J, Samulski RJ, Li C. 2018. Double-stranded RNA innate immune response activation from long-term adeno-associated virus vector transduction. Jci Insight 3:e120474.

48. Grimm D, Kern A, Rittner K, Kleinschmidt JA. 1998. Novel Tools for Production and Purification of Recombinant Adenoassociated Virus Vectors. Hum Gene Ther 9:2745–2760.

49. Grieger JC, Choi VW, Samulski RJ. 2006. Production and characterization of adeno-associated viral vectors. Nat Protoc 1:1412–1428.

50. Lawrence M, Huber W, Pagès H, Aboyoun P, Carlson M, Gentleman R, Morgan MT, Carey VJ. 2013. Software for computing and annotating genomic ranges. PLoS computational biology 9:e1003118.

51. Love MI, Huber W, Anders S. 2014. Moderated estimation of fold change and dispersion for RNA-seq data with DESeq2. Genome biology 15:550.

52. Luo W, Brouwer C. 2013. Pathview: an R/Bioconductor package for pathway-based data integration and visualization. Bioinformatics (Oxford, England) 29:1830– 1831.

53. Deschamps T, Kalamvoki M. 2017. Evasion of the STING DNA-Sensing Pathway by VP11/12 of Herpes Simplex Virus 1. J Virol 91.

54. Vandesompele J, Preter KD, Pattyn F, Poppe B, Roy NV, Paepe AD, Speleman F. 2002. Accurate normalization of real-time quantitative RT-PCR data by geometric averaging of multiple internal control genes. Genome biology 3:RESEARCH0034.

55. Sitaraman V, Hearing P, Ward CB, Gnatenko DV, Wimmer E, Mueller S, Skiena S, Bahou WF. 2011. Computationally designed adeno-associated virus (AAV) Rep 78 is efficiently maintained within an adenovirus vector. Proc Natl Acad Sci 108:14294–14299.

56. Kim E-J, Park J-I, Nelkin BD. 2005. IFI16 Is an Essential Mediator of Growth Inhibition, but Not Differentiation, Induced by the Leukemia Inhibitory Factor/JAK/STAT Pathway in Medullary Thyroid Carcinoma Cells*. J Biol Chem 280:4913–4920.

57. Herzner A-M, Hagmann CA, Goldeck M, Wolter S, Kübler K, Wittmann S, Gramberg T, Andreeva L, Hopfner K-P, Mertens C, Zillinger T, Jin T, Xiao TS, Bartok E, Coch C, Ackermann D, Hornung V, Ludwig J, Barchet W, Hartmann G, Schlee M. 2015. Sequence-specific activation of the DNA sensor cGAS by Y-form DNA structures as found in primary HIV-1 cDNA. Nat Immunol 16:1025–1033.

58. Lux K, Goerlitz N, Schlemminger S, Perabo L, Goldnau D, Endell J, Leike K, Kofler DM, Finke S, Hallek M, Büning H. 2005. Green fluorescent protein-tagged adeno-associated virus particles allow the study of cytosolic and nuclear trafficking. Journal of Virology 79:11776–11787.

